# The *Solanum americanum* pangenome and effectoromics reveal new resistance genes against potato late blight

**DOI:** 10.1101/2022.08.11.503608

**Authors:** Xiao Lin, Yuxin Jia, Robert Heal, Maxim Prokchorchik, Maria Sindalovskaya, Andrea Olave-Achury, Moffat Makechemu, Sebastian Fairhead, Azka Noureen, Jung Heo, Kamil Witek, Matthew Smoker, Jodie Taylor, Ram-Krishna Shrestha, Yoonyoung Lee, Chunzhi Zhang, Soon Ju Park, Kee Hoon Sohn, Sanwen Huang, Jonathan D. G. Jones

## Abstract

Late blight caused by the oomycete pathogen *Phytophthora infestans* continues to cause major worldwide losses in potato and tomato. Most accessions of *Solanum americanum*, a globally distributed, wild Solanaceae plant, are highly resistant to late blight. We generated high-quality reference genomes of four *S. americanum* accessions, re-sequenced 52 accessions, and we defined variation in the NLR immune receptor genes (the *S. americanum* NLRome). We further screened for variation in recognition of ∼315 *P. infestans* RXLR effectors in 52 *S. americanum* accessions. Using these genotypic and phenotypic data, we cloned three novel NLR-encoding genes *Rpi-amr4, Rpi-amr16* and *Rpi-amr17*, and determined their corresponding RXLR effector genes *Avramr4* (*PITG_22825*), *Avramr16* (*PITG_02860*) and *Avramr17* (*PITG_04373*) from *P. infestans*. These genomic resources and methodology will support efforts to convert potato into a “nonhost” of late blight and can be applied to diseases of other crops.

## Introduction

Potato is the most consumed non-grain crop worldwide. However, pests and diseases reduce global yields by ∼17% ^1^. Potato late blight, which is caused by the oomycete pathogen *Phytophthora infestans* ^2^, triggered the Irish famine in the 1840s, and is still the most damaging disease for global potato production ^1^.

Plant immunity depends on pathogen recognition by both cell-surface pattern recognition receptors (PRRs) and intracellular immune receptors. Most cloned plant *Resistance (R*) genes encode intracellular nucleotide-binding domain, leucine-rich repeat (NLR) proteins. Many *R* genes against *P. infestans* (*Rpi* genes) were cloned from wild relatives of potato species, such as *R2, R3, R8, Rpi-blb1, Rpi-blb2* and *Rpi-vnt1* from *Solanum demissum, S. bulbocastanum* and *S. venturii* ^3-10^. However, most cloned *Rpi* genes have been overcome by the fast-evolving pathogen.

NLR proteins recognize pathogen effectors, either directly or indirectly. *P. infestans* effectors carry a signal peptide and an RXLR (Arg-X-Leu-Arg, X represents any amino acid)-EER (Glu-Glu-Arg) motif. In the *P. infestans* reference genome (strain T30-4), 563 RXLR effectors were predicted, enabling screens for recognition of these effectors (“effectoromics”) in various plant genetic backgrounds ^11,12^.

Reference genome sequences of potato, tomato, eggplant and pepper have been determined ^13-16^. Phased, chromosome-level genome assemblies of heterozygous diploid and tetraploid potatoes are also available ^17-19^. Pan-genome studies of crop plants including potato have also emerged, that shed light on the extensive genetic variation in these species ^20-23^. To reduce the genomic complexity and sequencing costs, resistance gene enrichment sequencing (RenSeq) and RLP/K enrichment sequencing (RLP/KSeq) technologies were developed to sequence plant *NLR* and *PRR* gene repertoires ^24-26^. These methods led to many important applications such as AgRenSeq and the defining of the pan-NLRome of *Arabidopsis* ^27,28^.

Diploid *Solanum americanum* and its closely related hexaploid descendant *S. nigrum* are both highly resistant to LB. Previously, our group cloned *Rpi-amr1* and *Rpi-amr3* from different resistant *S. americanum* accessions, and their functional homologs were cloned from *S. nigrum*^25,29,30^. Both *Rpi-amr1* and *Rpi-amr3* confer late blight resistance in potato. The cognate effectors AVRamr1 and AVRamr3 were identified by pathogen enrichment sequencing (PenSeq), and effectoromics screening ^30,31^.

Here, we sequenced and assembled four high-quality genomes of *S. americanum* and re-sequenced 52 accessions that were collected from seed banks around the world. By manually annotated *NLR* genes we defined the Pan-NLRome of *S. americanum* in combination with additional 16 SMRT-RenSeq assemblies. We also screened 315 *P. infestans* RXLR effectors in 52 *S. americanum* accessions by transient expression through *Agrobacterium* infiltration. These genomic resources and functional data led to the rapid identification of three novel *Rpi* genes *Rpi-amr4, Rpi-amr16* and *Rpi-amr17* that are responsible for AVRamr4 (PITG_22825), AVRamr16 (PITG_02860) and AVRamr17 (PITG_04373) recognition. This study unveils a non-host effector-triggered immunity (ETI) interaction between *S. americanum* and *P. infestans*, that will enable us to clone more *Rpi* genes from the *S. americanum* gene pool. In combination with novel technologies for potato genome design and engineering ^32^, this will help develop potatoes with durable late blight resistance.

## Results

### Genome assembly and gene model prediction of *Solanum americanum*

*S. americanum* is a globally distributed Solanaceae species, which is resistant to many pathogens including *Phytophthora infestans* and *Ralstonia solanacearum*^25,29,33^. Four *S. americanum* accessions SP1102, SP2271, SP2273 and SP2275 were selected for sequencing based on their variation in resistance to late blight (Supplementary Fig. 1). We first generated Illumina paired-end reads to analyze these genomes. K-mer based analysis indicated that the genome sizes of these four *S. americanum* accessions were 1.15-1.31 Gb, with 0.05% to 0.35% heterozygosity (Table 1, Supplementary Fig. 2). We then generated an average of 29.5 Gb (∼26 fold) PacBio high-fidelity reads for SP1102 and SP2271, respectively, and assembled the reads into 298 and 568 contigs with N50s 82.9 Mb and 55.2 Mb. The ONT platform was used for sequencing SP2273 and SP2275 resulting in ∼81.1 Gb (∼62 fold) and ∼114.5 Gb ONT reads (∼95 fold), respectively. The noisy reads were self-corrected and assembled into 304 and 539 contigs with N50s 9.6 Mb and 4.8 Mb, respectively. To generate chromosome level assemblies, we further generated ∼86.5 Gb, ∼81.8 Gb and ∼54.8 Gb Hi-C data for SP1102, SP2271 and SP2273, and anchored the contigs into 12 pseudomolecules (Supplementary Fig. 3). The completeness of these assemblies was estimated to be ∼98.4% (single-copy and duplicated, Supplementary Fig. 4a) by BUSCO, which indicates the high quality of genome assembly.

**Table 1:**
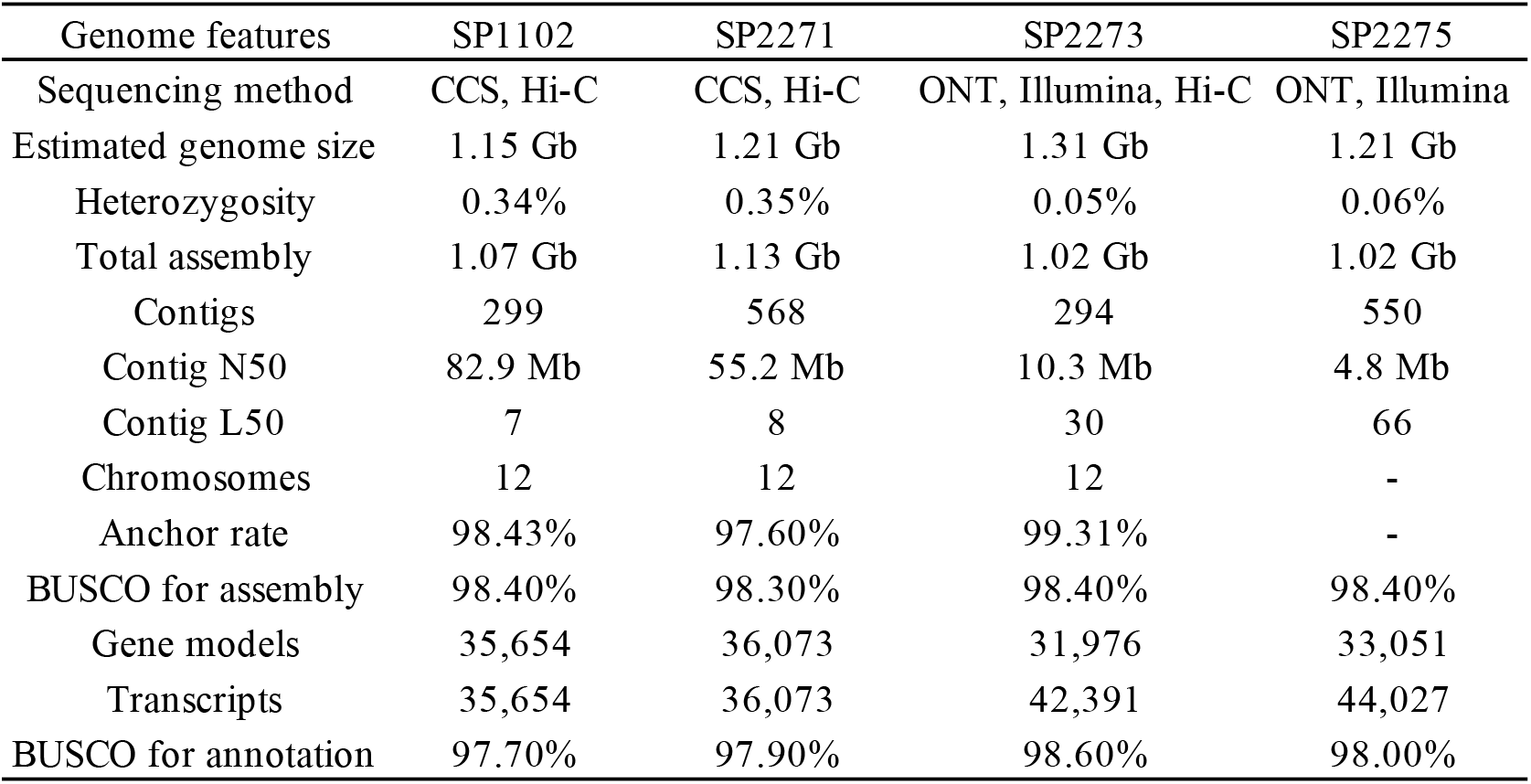

| Genome features | SP1102 | SP2271 | SP2273 | SP2275 |
| --- | --- | --- | --- | --- |
| Sequencing method | CCS, Hi-C | CCS, Hi-C | ONT, Illumina, Hi-C | ONT, Illumina |
| Estimated genome size | 1.15 Gb | 1.21 Gb | 1.31 Gb | 1.21 Gb |
| Heterozygosity | 0.34% | 0.35% | 0.05% | 0.06% |
| Total assembly | 1.07 Gb | 1.13 Gb | 1.02 Gb | 1.02 Gb |
| Contigs | 299 | 568 | 294 | 550 |
| Contig N50 | 82.9 Mb | 55.2 Mb | 10.3 Mb | 4.8 Mb |
| Contig L50 | 7 | 8 | 30 | 66 |
| Chromosomes | 12 | 12 | 12 | - |
| Anchor rate | 98.43% | 97.60% | 99.31% | - |
| BUSCO for assembly | 98.40% | 98.30% | 98.40% | 98.40% |
| Gene models | 35,654 | 36,073 | 31,976 | 33,051 |
| Transcripts | 35,654 | 36,073 | 42,391 | 44,027 |
| BUSCO for annotation | 97.70% | 97.90% | 98.60% | 98.00% |

**Figure 1.**
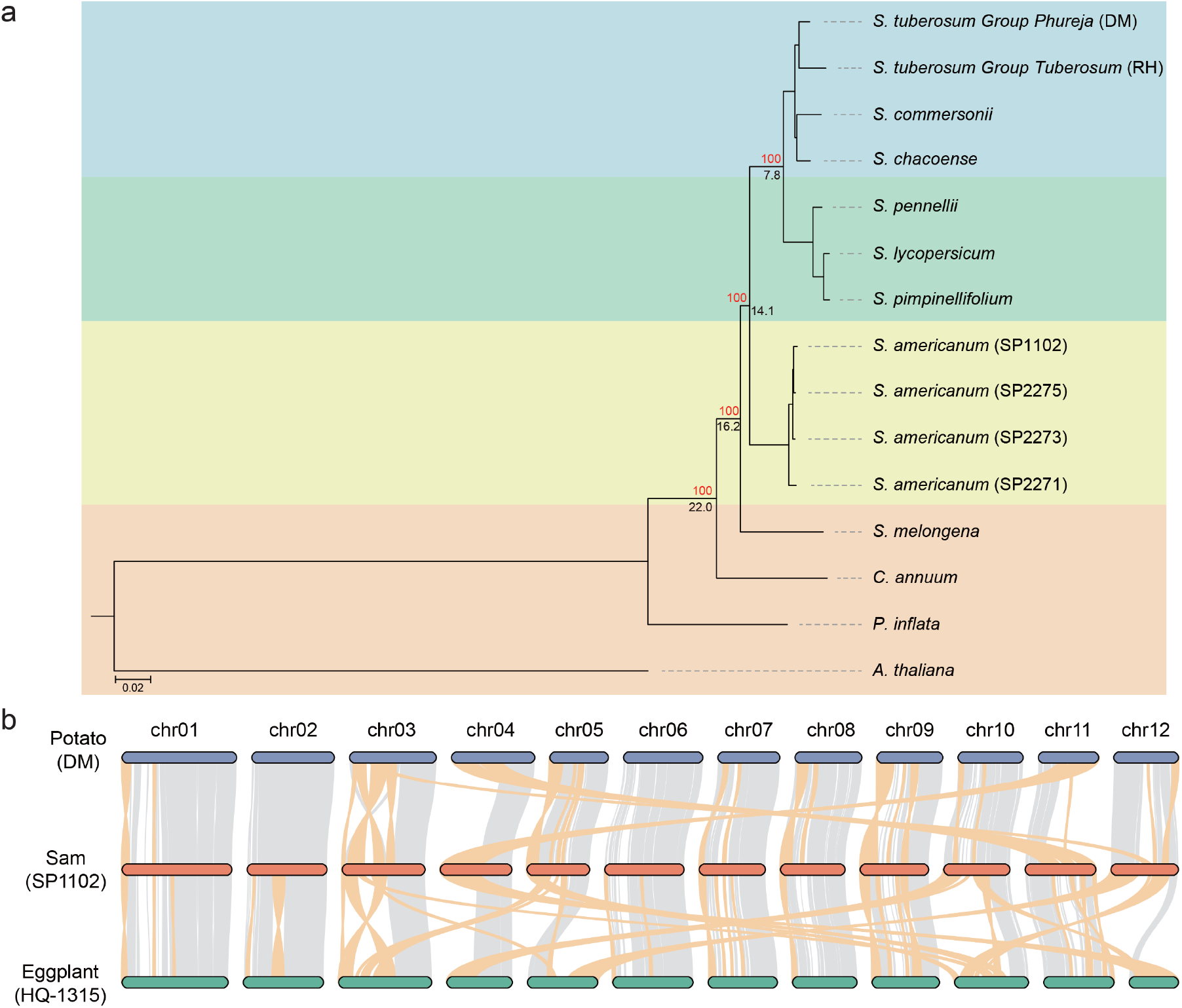
Genome evolution of *Solanum americanum*. (a) Phylogenetic relationship of *Solanum americanum* and neighbouring species. The red number indicates bootstrap of each node. The black number denotes estimated divergence time (million years ago). (b) Genome synteny of *Solanum americanum*, potato and eggplant. Ribbons between chromosomes show syntenic regions; Large chromosome rearrangements (> 1 Mb in size) are marked as orange.

**Figure 2.**
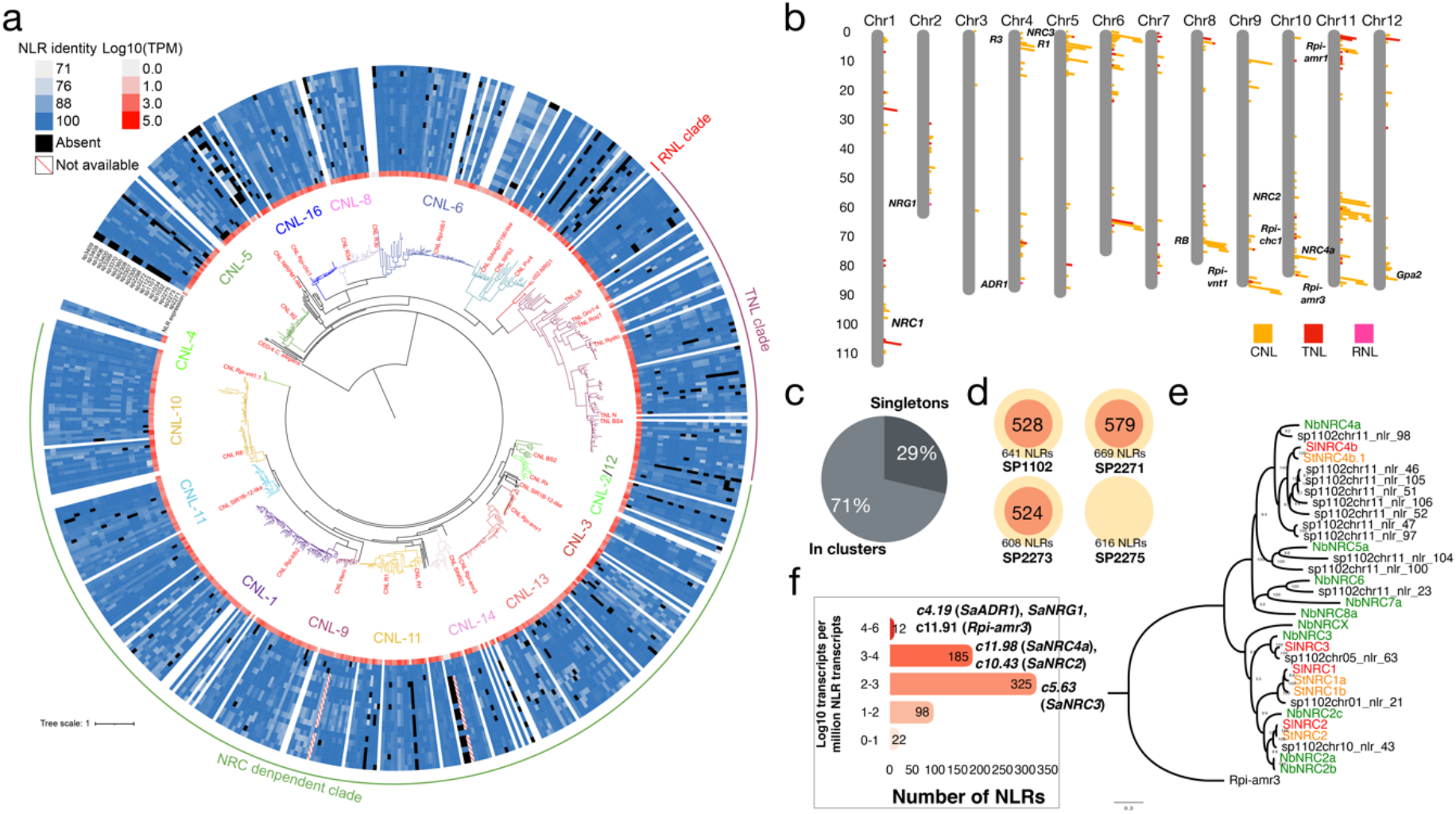
Pan-NLRome of *Solanum americanum*. (a) The NB-ARC domains of *S. americanum* SP1102 were predicted by NLR-annotator and used to generate the maximum-likelihood tree by IQ-TREE with JTT+F+R9 model. Known NLR proteins from Solanaceae species are included (highlighted by red). The NLRs are classified into different groups based on a previous report ^29^. The RNL, TNL and NRC-dependent CNL clade are shown. CED-4 from *Caenorhabditis elegans* were used as the outgroup. The expression profile is shown by a heatmap (white to red) based on the cDNA RenSeq data of SP1102. The P/A polymorphism of NLRs from the three other *S. americanum* genomes and SMRT-RenSeq assemblies of 16 additional accessions are shown by the heatmap (white to blue), the absent NLRs are shown by black blocks. (b) The physical map of *NLR* genes in SP1102 genome. CNLs are shown by yellow blocks, TNLs are red, and RNLs are pink blocks. Some functionally characterized *NLR* clusters are noted on this map. (c) The proportion of NLR singletons and NLRs in clusters. (d) Number of manually curated *NLR* genes, and the *NLR* number that predicted by NLR-annotator (manual curation of SP2275 was not performed). (e) Phylogeny of the NRC helper NLR family. The NRC homologs from potato, tomato and *N. benthamiana* are marked in yellow, red and green respectively. The *NRC* genes from *S. americanum* are black. (f) The log10 transcript per million NLR transcripts are classified into 5 groups, and some homologs of known *R* genes are noted. The NLR IDs are shown in Supplementary Table 1.

**Figure 3.**
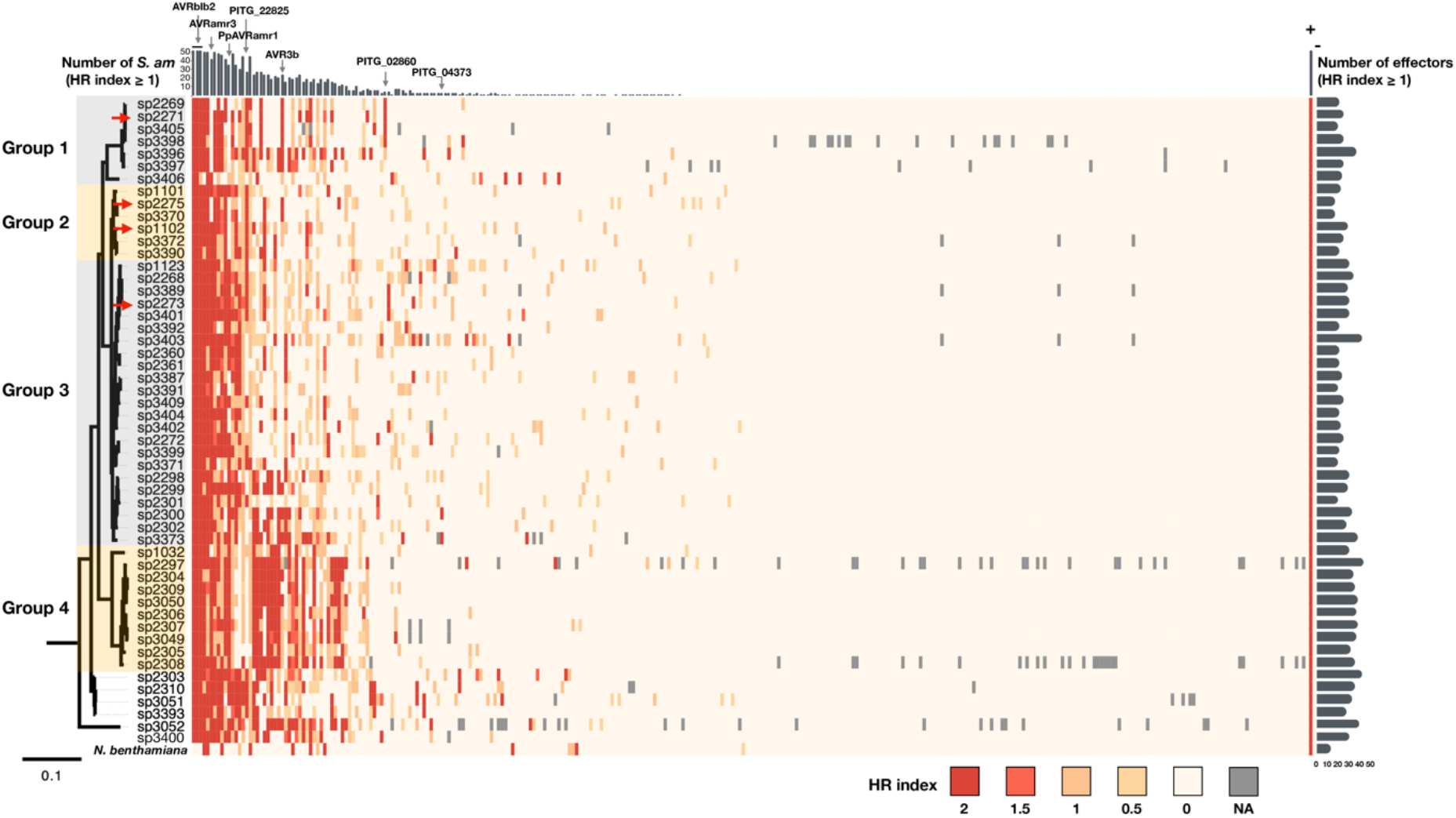
ETI landscape of *S. americanum* and *P. infestans*. 315 RXLR effectors were transiently expressed in 52 *S. americanum* accessions. The HR index (strong HR = 2, weak HR = 1, no HR = 0, NA = not available) was used for the heatmap. These effectors were screened on *N. benthamiana*^30^, their recognitions are included in this heatmap. Empty PVX vector was used as negative control, co-expression of Rpi-amr3::HF and AVRamr3::GFP was used as positive control. The *S. americanum* accessions were ordered based on the phylogenetic tree, SP3400 is not included in this tree. The *S. americanum* accessions were classified into 4 groups (grey or yellow shading). The 4 reference accessions are marked by red arrows. The effectors were ordered based on the total HR index. For each effector, the numbers of responsive *S. americanum* accessions are visualized by a bar chart on the top of the heatmap. For each *S. americanum* accession, the numbers of recognized effectors are visualized by a bar chart on the right of the heatmap. Some RXLR effectors previously characterised or mentioned in this study are indicated by grey arrows.

**Figure 4.**
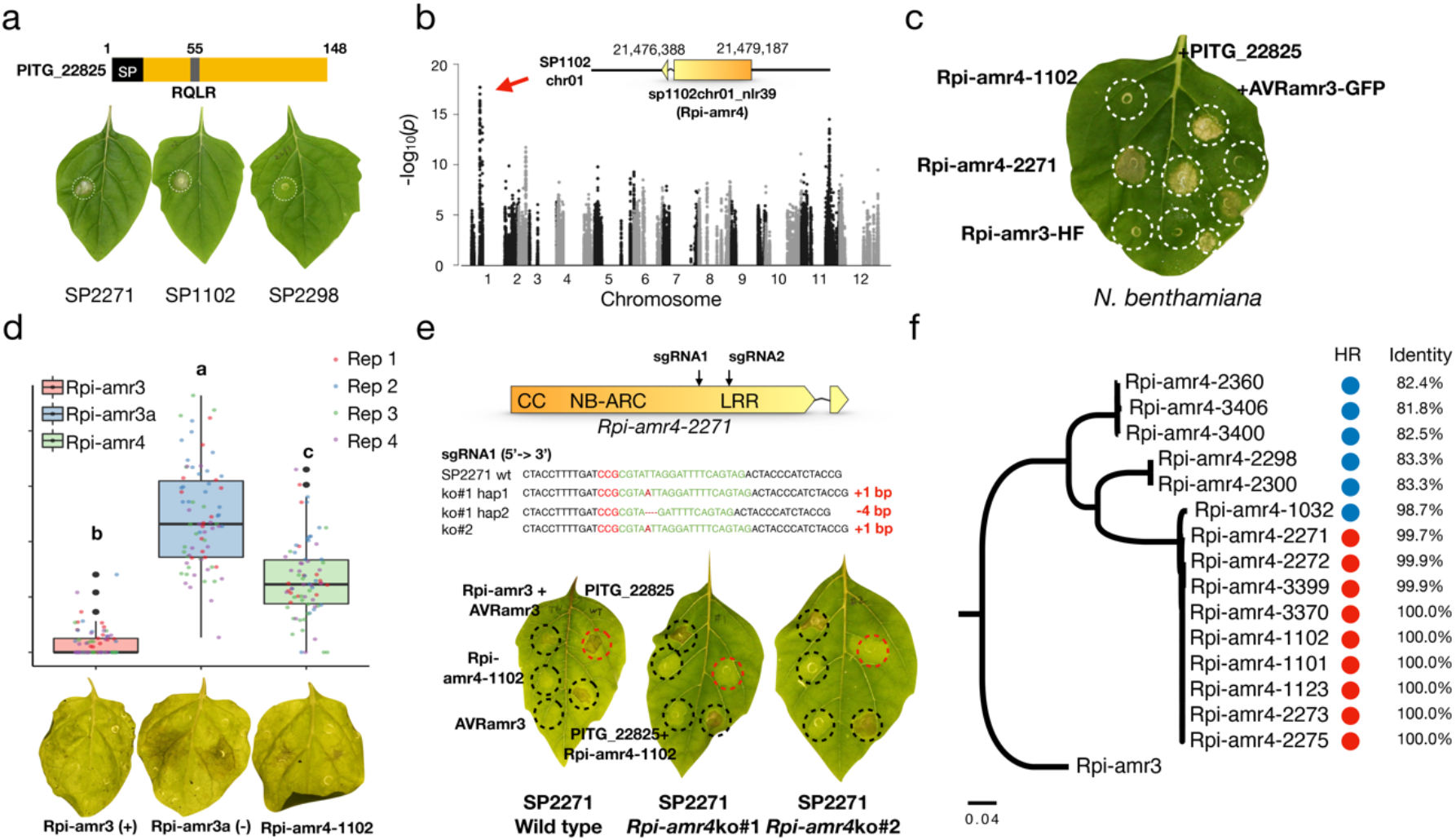
Identification and characterisation of *Rpi-amr4* that recognizes PITG_22825. (a) PITG_22825 is an RXLR effector with a signal peptide (SP), an RQLR-EER motif and an effector domain. The effector without SP was cloned into a vector under control of 35S promoter and transformed into *Agrobacterium tumefaciens* strain GV3101-pMP90 for transient expression through agroinfiltration, and triggers cell death on *S. americanum* SP2271 and SP1102, but not SP2298. (b) Manhattan plot of the GWAS of PITG_22825 recognition. Resequencing reads from 52 *S. am* accessions were mapped to the *S. americanum* SP1102 genome, the SNPs on *NLR* genes were called and filtered for the GWAS. SNPs significantly associated with PITG_22825 recognition are located on the NLR singleton sp1102chr01_nlr39 (red arrow). (c) HR assay of candidate genes. *Rpi-amr4-1102* and *Rpi-amr4-2271* were cloned into a vector under control of 35S promoter, then transformed into *Agrobacterium tumefaciens* strain GV3101-pMP90. They were expressed alone, or co-expressed with either 35S::PITG_22825 or 35S::AVRamr3-GFP constructs in *N. benthamiana* leaves. Rpi-amr4-2271 is auto-active in *N. benthamiana*, but when co-expressed with PITG_22825, the HR was stronger. Rpi-amr4-1102 specifically recognize PITG_22825 but not AVRamr3. Rpi-amr3 was used as a control. OD_600_ = 0.5. Four leaves from two plants were used for each experiment, and three biological replicates were performed with same results. (d) Detached leaf assay (DLA) of *Rpi-amr4. Rpi-amr4* (green), *Rpi-amr3* (positive control, red) and *Rpi-amr3a* (a non-functional *Rpi-amr3* paralogue, negative control, blue) were transiently expressed in *N. benthamiana*, OD_600_ = 0.5. The zoospores (300 zoospores in a 10 µL droplet) from *P. infestans* strain T30-4 were used to inoculate the leaves 1 day after the infiltration. A calliper was used to measure the lesion size at 6 days after inoculation. Four biological replicates were performed, all datapoints (74 datapoints/treatment) were visualized as a box plot using R. The outliers are indicated by black dots. Statistical differences were analysed by one-way ANOVA with Tukey’s HSD test (P < 0.001). The representative leaves of this assay are shown. (e) *Rpi-amr4* knockout lines lose the PITG_22825 recognition capacity. Two sgRNAs (black arrows) were designed to knockout *Rpi-amr4*-2271 in SP2271. The genotype of two knockout lines is shown. The two knockout lines failed to recognize PITG_22825, but the HR could be complemented by co-expressed PITG_22825 with *Rpi-amr4-*1102. Wild-type SP2271 was used as control plants. *Rpi-amr3* and AVRamr3 were used as control for the agroinfiltration. OD_600_ = 0.5, four-weeks SP2271 plants were used for agroinfiltration, the photos were taken 4 days after infiltration. (f) Phylogeny of Rpi-amr4 homologs in different *S. americanum* accessions. Rpi-amr3 was used as an outgroup. The PITG_22825-mediated HR are shown by red (HR) or blue (no HR) circles. Percent identity of amino acid sequence relative to Rpi-amr4-1102 is shown. The scale bars represent the number of amino acid substitutions per site.

To annotate gene models of SP1102 and SP2271, we generated an average of ∼80 Gb transcriptome data (150 bp, paired-end reads) for each accession, which derived from young seedlings, roots, stems, leaves, flowers and fruits. The EVidenceModeler pipeline was applied to annotate the gene models by integrating *ab initio* prediction, homology-based annotation and transcriptome evidence. To predict gene models in SP2273 and SP2275 genomes, we deployed the GeMoMa pipeline to integrate homology-based annotation and transcriptome evidence. In summary, we predicted an average of 34,193 gene models with an average of 98.1% (single-copy and duplicated, Supplementary Fig. 4b, Table 1) BUSCO evaluation for each *S. americanum* genome.

### Genome evolution of *S. americanum*

To investigate the evolution of *S. americanum* genomes, we clustered the representative protein sequences from 15 genomes, comprising four *S. americanum*, four potatoes, three tomatoes, four additional Solanaceae species and an outgroup species of *Arabidopsis thaliana*, into 33,115 orthogroups, from which we further identified 1,363 single-copy orthogroups. Using a supermatrix method, we inferred the phylogenetic relationships and estimated the divergence time of the 15 genomes. The species tree topology suggests that *S. americanum* is a sister species to the common ancestor of potato and tomato, and they diverged ∼14.1 million years ago (Ma, 95% highest posterior density interval: 11.7-17.2 Ma, Fig. 1a), which is consistent with the former report based on plastid sequences ^34^.

Chromosome rearrangement (CR) is an important evolutionary process ^35^. The reference-grade genome assemblies enabled us to explore *S. americanum* chromosome evolution. Both protein coding gene-based chromosome syntenic analysis and alignment based on genomic sequences were carried out and generated similar results (Fig. 1b, Supplementary Fig. 5). We observed 45 large CRs (> 1 Mb in size), comprising 26 inversion and 19 inter-chromosome translocation events, between *S. americanum* and potato genomes (Fig. 1b). In contrast, 67 large CRs (30 inversions and 37 inter-chromosome translocations) were found between *S. americanum* and eggplant (Fig. 1b). Notably, CRs were not evenly distributed across the genome. No CR was identified on chromosome 2 between *S. americanum* and potato, while as many as 11 CRs occurred on chromosome 11.

**Figure 5.**
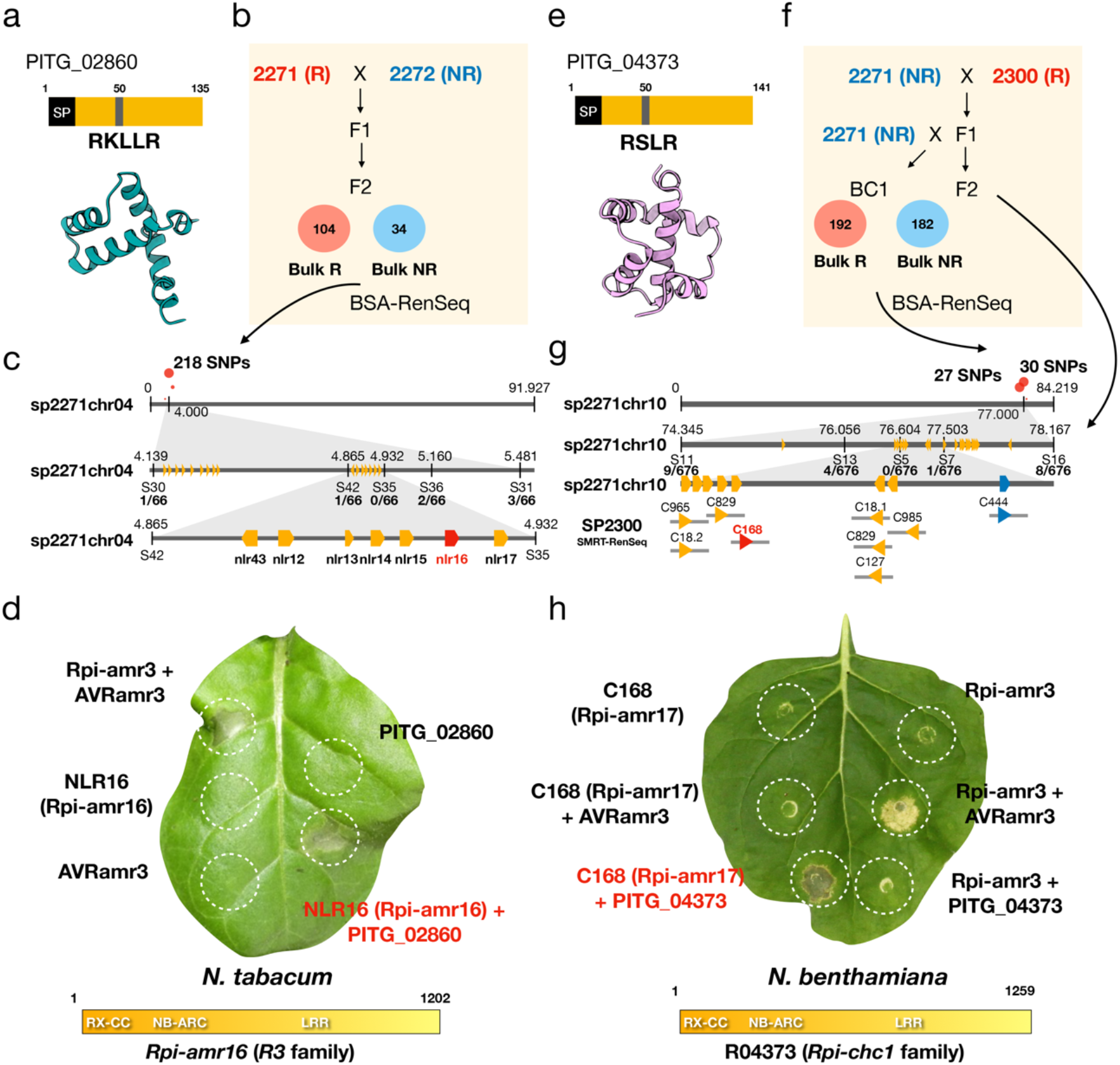
Identification of *Rpi-amr16* and *Rpi-amr17* that recognize PITG_02860 and PITG_04373. (a) PITG_02860 is an RXLR effector from *P. infestans*, an illustration and AlphaFold predicted structure are shown. (b) An F2 population was generated from a cross between SP2271 (Responds to PITG_02860, R) and SP2272 (No response to PITG_02860, NR). Within this population PITG_02860 responsiveness segregates in a ratio indicating a single dominant gene is responsible. The responsive bulk (Bulk R) and non-responsive bulk (Bulk NR) were used for BSA-RenSeq. (c) The BSA-RenSeq analysis led to the identification of 218 informative SNPs (red dots) on the top of Chr04 of SP2271. The grey bar represents the chromosome. The physical positions (MB) are shown by number. Five molecular markers (S30, S42, S35, S36 and S31) were designed and used for the map-based cloning. Number of recombination events/total tested gametes are shown below the marker name. the physical positions are indicated in Mb. (d) HR assay of the candidate PITG_02860 receptor. The candidates were cloned into a vector with 35S promoter and transformed into *Agrobacterium tumefaciens* strain GV3101-pMP90. They were expressed alone, or co-expressed with PVX::PITG_02860 in *N. tabacum* leaves, Rpi-amr3-HF and AVRamr3-GFP were used as controls. NLR16 turned out to be the PITG_02860 receptor (*Rpi-amr16* hereafter). OD_600_ = 0.5, four-week *N. tabacum* plants were used for agroinfiltration, the photos were taken at 4 days after infiltration. Four leaves from two plants were used for each experiment, and three biological replicates were performed with same results. *Rpi-amr16* encodes a 1202 aa protein, and it belongs to the *R3* family. (e) PITG_04373 is an RXLR effector from *P. infestans*, the illustration and predicted structure are shown. (f) Both backcross (BC1) and F2 populations were generated from SP2271 and SP2300. The BC1 population of 192 responsive plants and 182 non-responsive progenies were bulked for BSA_RenSeq. The F2 population were used for fine mapping. (g) BSA-RenSeq led to the identification informative SNPs (red dots) on the top of Chr10 of SP2271. The grey bar represents the chromosome. The physical positions (MB) are shown by number. Five molecular markers (S11, S13, S5, S7 and S16) were designed and used for fine mapping. Number of recombination gametes/ total tested gametes are shown below the marker name. SMRT_RenSeq assemblies of SP2300 were mapped to SP2271 genome, and nine genes which mapped to the mapping interval were identified as candidate genes of PITG_04373 receptor. *C444* belongs to *R1* family, all other candidate genes belong to *Rpi-chc1* family. (h) HR assay of the PITG_04373 receptor candidates. Genes were cloned into a vector with 35S promoter and transformed into *Agrobacterium tumefaciens* strain GV3101-pMP90. They were expressed alone, or co-expressed with 35S::PITG_04373 in *N. benthamiana* leaves, Rpi-amr3-HF and AVRamr3-GFP were used as controls. C168 turned out to be the PITG_04373 receptor (Rpi-amr17 hereafter). OD_600_ = 0.5, four-weeks *N. benthamiana* plants were used for agroinfiltration, photos were taken at 4 days after infiltration. Four leaves from two plants were used for each experiment, and three biological replicates were performed with same results. *Rpi-amr17* encodes a 1259 aa protein, and it belongs to the *Rpi-chc1* family.

### **Structural variation between** *S. americanum* **genomes**

Structural variations (SVs), including insertions, deletions, duplications, inversions and translocations, cause and maintain phenotypic diversity ^36^. The chromosome assemblies of three *S. americanum* genomes enabled analysis of large SVs (> 1 Mb in size). Using SP1102 as the reference, we identified 56 large SVs in SP2271 (Supplementary Fig. 6), impacting ∼256 Mb of the reference genome. However, only 14 large SVs were identified in SP2273, covering ∼54 Mb of the reference genome (Supplementary Fig. 6). The huge differences of SV number between *S. americanum* genomes shed light on the complex history of *S. americanum* species evolution.

To further characterize SVs within *S. americanum*, we used the SP1102 as the reference and identified 60,849, 23,672 and 18,492 SVs (>40 bp in size) in SP2271, SP2273 and SP2275, ranging from 27.9 to 102.0 Mb in total length (Supplementary Fig. 7). Most SVs (∼54.5% on average) located in putative gene regulatory regions (5 kb upstream, downstream or in introns), followed by 44.0% and 1.5% overlapped with intergenic region and exon. SVs might contribute to gene expression variation. We identified 1,837 differentially expressed genes (DEGs, Fold change >= 2 or <= 0.5, P-value < 0.05) in leaves between SP1102 and SP2271, of which, 1,084 DEGs might associate with SVs, as exemplified by *sp1102chr11_nlr_6*, an NLR gene whose expression might be impaired by a 286 bp deletion in the promoter region in leaves of SP2271 (Supplementary Fig. 8).

Previous reports indicate that NLR genes in plants are highly diversified between individuals even in the same species ^37^. We found that sequence diversity in NLR region is significantly higher than that of non-NLR region (Supplementary Fig. 9a). As an example, the *Rpi-amr3* locus varied greatly among *S. americanum* genomes (Supplementary Fig. 9b), suggesting the NLRome based on high-quality genomes is required for deep understanding the non-host resistance of *S. americanum* species.

### Defining the *S. americanum* pan-NLR repertoire

To understand *NLR* gene diversity, a phylogenetic tree was generated by using the NB-ARC domain of the NLR proteins from SP1102 (Fig. 2a), and the physical positions of these *NLRs* in SP1102 genome were visualized in the physical map (Fig. 2b). We found 71% *NLRs* in the SP1102 genome are in clusters and the rest are singletons (Fig. 2c). Due to the complexity of *NLR* gene clusters, most automatic annotation pipelines produce incorrect gene models^37^. To generate better gene models of the *NLR* genes from *S. americanum* genome. Here, we first used NLR annotator ^38^ to identify the putative *NLR* genes from SP1102, SP2271, SP2273 and SP2275 reference genomes, revealing 641, 669, 608 and 616 *NLRs* respectively (Fig. 2d). By adopting a definition of *NLR* cluster (3 *NLRs* within 200 kb interval) ^39^. To predict the precise *NLR* gene models, we manually annotated 528, 579 and 524 *NLRs* from SP1102, SP2271 and SP2273 genomes by incorporating NLR-annotator results and cDNA sequence data (Fig. 2d, Supplementary dataset 1-3). Next, we examined presence/absence (P/A) polymorphism of NLRs among *S. americanum* accessions. NLRs from 3 other *S. americanum* reference genomes, and from 16 SMRT-RenSeq assemblies, were compared with the SP1102 NLRs. P/A polymorphism is shown in Fig. 2a. We also inspected the expression level of SP1102 NLRs by re-analysing cDNA RenSeq data ^25^. The transcripts per million *NLR* transcripts were visualized by a heatmap (Fig. 2a). Many well-known *NLR* genes are relatively highly expressed, such as *ADR1, NRG1, Rpi-amr3, NRC4a, NRC2* and *NRC3* (Fig. 2f). Many Solanaceae CNLs require NRC helpers to execute their function and 50% of the *S. americanum* NLRs lie within the NRC superclade ^40^ (Fig. 2a). To investigate the NRC family, we generated another phylogenetic tree for NRCs. We found NRC1, NRC2, NRC3, NRC4a and NRC6 homologs, two NRC5a copies. Interestingly, the copy number of NRC4b homologs has expanded in the *S. americanum* genome (Fig. 2e). Rpi-amr3 and Rpi-amr1 require NRC2/3/4 and NRC2/3, respectively ^29,30^. However, *NRC1* is missing in *N. benthamiana* ^41^. To test if NRC1 can enable *Rpi-amr1/3* function, we cloned the *SaNRC1* from SP1102 and showed *SaNRC1* enables *Rpi-amr3* but not *Rpi-amr1* function in *N. benthamiana* nrc2/3/4 knockout line (Supplementary Fig. 10). In summary, we generated an NLRome of ∼20 *S. americanum* accessions, and manually annotated the *NLRs* from 3 reference genomes. This resource is important for the investigation *NLR* gene evolution, and functional studies of ETI in *S. americanum* and other Solanaceae species.

### The ETI landscape of the *S. americanum* and *P. infestans* interaction

There are 563 predicted RXLR effectors from *P. infestans* reference genome T30-4. In this study, we showed that there are ∼550 *NLR* genes in the best-studied *S. americanum* reference genomes (Fig. 2d). To reveal one-to-one effector-receptor interactions and clone more *Rpi* genes, we screened ∼315 RXLR effectors on 52 *S. americanum* accessions (Fig. 3). Based on the cDNA PenSeq data, most if not all expressed RXLR effectors were included in this screening ^31^. We found that 5 effectors triggered HR on most *S. americanum* accessions (≥ 50), including effectors from AVRblb2 family. 185 effectors did not trigger HR in any *S. americanum* accessions, 71 effectors can only be recognized by less than 5 *S. americanum* accessions. Notably, the rest 54 effectors showed differential recognition by different *S. americanum* accessions. AVRamr1 and AVRamr3 are also widely recognized by different *S. americanum* accessions (Fig. 3). The 4 reference accessions SP2271, SP2275, SP1102 and SP2273 can recognize 25, 18, 30 and 30 RXLR effectors respectively, of which 5, 3, 7 and 9 effectors are specifically recognized in each accession (Supplementary Fig. 11). Notably, accession SP2271 is susceptible to *P. infestans* in the detached leaf assay (DLA), yet resistant to late blight in the field. As expected, SP2271 does not recognize AVRamr1 and AVRamr3. We found premature stop codons in both *Rpi-amr1* and *Rpi-amr3* SP2271 homologs. Intriguingly, 25 RXLR effectors trigger HR in SP2271, these RXLR effector recognitions might contribute to the SP2271 field resistance to late blight. Conceivably, effector recognitions from other *S. americanum* accessions other than SP2271 contribute to stronger resistance against late blight. These results reveal the ETI landscape of *S. americanum* against the late blight pathogen and enable identification of novel *Rpi* genes.

### Resequencing of *S. americanum* accessions

To understand the genetic polymorphism of all *S. americanum* accessions in our collection, we performed PCR-free, 150 bp paired-end sequencing for 52 *S. americanum* accessions at 10x coverage. The reads were mapped to the reference *S. americanum* genome SP1102, and SNPs were called and filtered for further analysis. We constructed a phylogenetic tree using all SNPs on genes. Eggplant, potato, and tomato were used as outgroups (Supplementary Fig. 12a). The structural and inbreeding coefficient values were also analysed (Supplementary Fig. 12b and c). The 52 accessions can be clearly assigned into 4 groups (Supplementary Fig. 12a). Group 1 contains SP2271, which is susceptible to late blight in lab conditions. All the 6 members in this group lack *Rpi-amr1* and *Rpi-amr3* based on the effectoromics screening (Fig. 3 and Supplementary Table 2). SP2275 and SP1102 are in Group 2, SP2273 is in Group 3. For Group 4, no reference genome is available, but we have SMRT RenSeq assemblies for several of these accessions. Surprisingly, four accessions SP2303, SP2310, SP3393 and SP3051 are not closely related to other groups and are highly heterozygous (Supplementary Fig. 12c), suggesting they may be polyploid species like *S. nigrum*. Two other accessions, SP3052 and SP3376 are also not closely related to the four *S. americanum* groups and might belong to another Solanaceae species. In summary, we re-sequenced 52 accessions and reveal their phylogenetic relationship and population structure. These data could be used for genome-wide association studies (GWAS), and molecular marker development.

### Cloning of *Rpi-amr4* by GWAS and linkage analysis

In our effectoromics screening, we found an effector PITG_22825 that triggers HR in 28/50 *S. americanum* accessions (Fig. 3, Supplementary Table 2), including SP1102 and SP2271 but not SP2298 (Fig. 4a). PITG_22825 is an RXLR effector with a signal peptide, RQLR and EER motifs followed by the effector domain (Fig. 4a). This effector had not received attention prior to our cDNA-PenSeq study ^31^. To map the gene conferring its recognition, a GWAS analysis was performed to associate this effector responsiveness with linked SNPs. A strong signal was identified on an NLR singleton that locates on Chr01 of SP1102 (Fig. 4b, Fig. 2b). This encodes a CNL that belongs to the Rpi-amr3 phylogenetic clade CNL-13 (Fig. 2a), but not physically in the Rpi-amr3 gene cluster on Chr11 (Fig. 2b). This likely explains another weaker GWAS signal on the *Rpi-amr3* cluster of Chr11 (Fig 2b). Based on cDNA RenSeq data from SP1102, the corresponding *NLR* gene carries an extra exon compared to *Rpi-amr3* (Fig. 4b). To verify this GWAS signal, we performed a bulked segregant analysis and resistance gene enrichment sequencing (BSA-RenSeq) in a segregating F2 population of SP2298 x SP2271 (Supplementary Fig. 13). The PITG_22825 responsive gene from SP2271 was mapped to the same position.

To test the gene function, the ORF of the candidate genes from SP2271 and SP1102 were PCR amplified and cloned into an over-expression binary vector with 35S promoter and Ocs terminator and introduced into *Agrobacterium*. The candidate genes were heterologously expressed alone, or co-expressed with PITG_22825 or AVRamr3-GFP (negative control) in *N. benthamiana*. Rpi-amr3-FLAG was used as a positive control. We found that the SP2271 allele (*Rpi-amr4-2271* hereafter) is auto-active in *N. benthamiana*, but when co-expressed with PITG_22825, the HR was faster and stronger compared to the control (Fig. 4c). In contrast, the SP1102 allele (*Rpi-amr4-1102* hereafter) is not auto-active in *N. benthamiana*, HR was triggered when *Rpi-amr4-1102* was co-expressed with PITG_22825, but not with AVRamr3-GFP (Fig. 4c). Rpi-amr3 does recognize AVRamr3-GFP but not recognize PITG_22825 but does recognize AVRamr3-GFP (Fig. 4c). There are only 3 amino-acid differences between *Rpi-amr4-1102* and *Rpi-amr4-2271*, and these differences might cause the auto-activity of the SP2271 allele (Supplementary Fig. 14). We also found *Rpi-amr4* is conserved in the PITG_22825 responsive accessions (Fig. 4f). To verify the function of *Rpi-amr4*, we designed two specific CRISPR-Cas9 sgRNAs and mutated *Rpi-amr4-2271* in SP2271. In total, sixteen CRISPR-Cas9 knockout lines were generated (Fig. 4e and Supplementary Table 4). The genotype and phenotype of two lines are shown in Fig 4e. Wild-type SP2271 can recognize PITG_22825 but the *Rpi-amr4*-mutated lines cannot. The HR phenotype could be complemented when *Rpi-amr4-1102* was co-expressed with PITG_22825 in the knockout lines (Fig. 4e). Therefore, we conclude that *Rpi-amr4* encodes the PITG_22825-recognizing immune receptor, and that *PITG_22825* is *Avramr4*.

To test if *Rpi-amr4* confers late blight resistance, we transiently expressed *Rpi-amr4-1102* in *N. benthamiana* leaves and inoculated with *P. infestans* zoospores. *Rpi-amr3* was used as a positive control and non-functional *Rpi-amr3a* from SP1102 was used as a negative control (Fig. 4d). This assay shows that *Rpi-amr4-1102* confers resistance against *P. infestans* isolate T30-4. However, the resistance is weaker compared to Rpi-amr3 (Fig. 4d). We also generated stable *Rpi-amr4-1102* transgenic *N. benthamiana* lines. As expected, the T0 transgenic plants gain the PITG_22825 recognition capacity, and they are resistant to two *P. infestans* isolates T30-4 and 88069 (Supplementary Fig. 15).

In summary, we successfully cloned a novel *Rpi* gene *Rpi-amr4* from *S. americanum* and defined its cognate effector gene *Avramr4* (*PITG_22825*). *Rpi-amr4* confers late blight resistance and may serve as resource for producing late blight resistant potatoes.

### Cloning of *Rpi-amr16* and *Rpi-amr17*

Although *Rpi-amr4* could be identified using GWAS approach, those effectors only recognized by a few *S. americanum* accessions did not enable a clear GWAS signal. We therefore deployed BSA-RenSeq for cloning two such effector responsive genes.

PITG_02860 (Fig 5a) targets a host protein NRL1, attenuates plant immunity and increases pathogen virulence ^42^. We found PITG_02860 triggers HR in 5/52 *S. americanum* accessions, including SP2271. We tested the F2 population of SP2271 (PITG_02860 responsive, R) x SP2272 (PITG_02860 non-responsive, NR), and found the recognition of PITG_02860 segregated according to a 3:1 ratio (104 R and 34 NR, χ^2^ (1, *N* = 138) = 0.00966, *p* = 0.92169) (Fig. 5b). gDNA was isolated from the R and NR bulks respectively, RenSeq libraries were constructed and sequenced (250 bp paired-end reads). The reads were mapped to SP2271, and SNPs were called and filtered. Most filtered SNPs located within a 1 Mb region on the top of Chr04 of SP2271 (Fig. 5c). To develop molecular markers for map-based cloning. The whole genome sequencing reads of SP2272 were mapped to SP2271 genome, then SCAR markers were designed to specifically amplify the responsive allele from SP2271, and the 34 NR progenies were then used for genotyping. The candidate gene was mapped to a 295 kb region between markers S42 and S36. Seven *NLRs* reside within the mapping interval, all from the *R3* family (Fig. 5c, Fig. 2a and b). To test these candidate genes, the ORFs from 4 candidate genes (*nlr13, nlr14, nlr16, nlr17*) were cloned into a binary vector under control of 35S promoter and Ocs terminator and transformed into *Agrobacterium* for transient expression. The candidate genes were expressed alone or with PITG_02860 or AVRamr3 in *N. benthamiana* and *N. tabacum*. NLR16 and NLR17 are auto-active in *N. benthamian*, but we found co-expression of NLR16 and PITG_02860 activate HR in *N. tabacum* (Fig. 5d). To verify this finding, we generated nlr16 knockout SP2271 lines by CRISPR-Cas9 system. As expected, the knockout lines lost recognition of PITG_02860 (Supplementary Fig. 16). Therefore, we conclude NLR16 (Rpi-amr16 hereafter) is the immune receptor for PITG_02860 (AVRamr16).

A sequence corresponding to PITG_04373 was designed and synthesized based on the PacBio and cDNA pathogen enrichment sequencing (PenSeq) data ^31^ (Fig. 5e). PITG_04373 triggers HR in only 3/50 *S. americanum* accessions including SP2300, which carries both functional *Rpi-amr1* and *Rpi-amr3* (Fig. 3). To clone the corresponding *Rpi* gene, we first phenotyped a BC1 population of SP2271(NR) x SP2300 (R) by transiently expressed PITG_04373 by agroinfiltration. The BC1 population segregates for PITG_04373 responsiveness with a 1:1 ratio (198 R : 182 NR, χ^2^ (1, *N* = 380) = 0.67368, *p* = 0.41177) (Fig. 5f). The DNA from R or NR progenies were bulked for BSA-RenSeq and most linked SNPs mapped to SP2271 Chr10. SCAR markers were designed by the same strategy and the candidate gene was ultimately mapped to a 1.447 Mb interval with 8 genes in SP2271 (Fig. 5g). Most candidates belong to the *Rpi-chc1* family, except a *R1* homolog (Fig. 5g). In the absence of a reference genome for SP2300, we generated a SMRT RenSeq assembly for the SP2300 NLRome ^29^. The SMRT-RenSeq contigs were mapped to this region of the SP2271 genome, and candidate genes from SP2300 were cloned into a vector with 35S promoter and Ocs terminator for transient assays. Five candidate genes were tested (C18.1, C18.2, C127, C168 and C829) by agroinfiltration with or without the effector. We found that the candidate gene C168 (Rpi-amr17 hereafter) can specifically recognize PITG_04373 after transient expression in *N. benthamiana* (Fig. 5h). Therefore, we conclude that *Rpi-amr17* encodes the PITG_04373 immune receptor from SP2300, and that PITG_04373 is AVRamr17.

To verify the function of *Rpi-amr17* in SP2300, we designed two sgRNAs targeting *Rpi-amr17* (Supplementary Fig. 17d). SP2300 also carries functional *Rpi-amr1* and *Rpi-amr3* homologs, to test if SP2300 carries additional *Rpi* genes, we also designed one sgRNAs for *Rpi-amr1-2300* and *Rpi-amr3*, respectively (Supplementary Fig. 17b and c). Forty transgenic SP2300 knockout lines were generated and tested by the three effectors (PpAVRamr1, AVRamr3 and PITG_04373), and we found 13 lines that lack the recognition for all three effectors; two such lines are shown (Supplementary Fig. 17e). We genotyped 2 knockout lines SLJ25603#3 and SLJ25603#17, and confirmed that all *Rpi-amr1-2300, Rpi-amr3-2300* and *Rpi-amr17* were knocked out (Supplementary Fig. 1b-d). We also co-expressed *Rpi-amr17* with *PITG_04373*, however the HR phenotype was not restored in this genotype. Interestingly, the triple knockout lines show some susceptibility to *P. infestans* (Supplementary Fig. 17f) compared to the wild-type SP2300, but it is still more resistant compared to SP2271, suggesting that there are additional *Rpi* genes in SP2300.

In summary, by using BSA-RenSeq, SMRT-RenSeq and map-based cloning strategies, we successfully cloned two novel *Rpi* genes *Rpi-amr16* and *Rpi-amr17* that confer recognition of two RXLR effectors PITG_02860 and PITG_04373.

## Discussion

*Solanum* is the largest genus of the Solanaceae family, comprising more than 1500 species, including many important crop plants such as potato, tomato and eggplant for which extensive genome sequence data are available. However, less is known about wild *Solanum* species. The *Solanum nigrum* complex is composed of many species with different ploidy levels, including *S. nigrum* (6x), *S. scabrum* (6x), *S. villosum* (4x) and *S. americanum* (2x). Some are regarded as weeds but others are consumed as food and medicine in various countries ^43^. Importantly, these species carry valuable natural sources of genetic resistance to diseases such as potato late blight and bacterial wilt ^25,29,33^. Variation for resistance to other pests and diseases is being investigated but when defining corresponding genes, reference genomes are highly advantageous. In this study, we sequenced and assembled four *S. americanum* genomes, three with Hi-C data that enable chromosome level assembly. These assemblies of *S. americanum* enabled us to gain insights into the interspecies structural variation, genomic evolution, and speciation of cultivated and wild Solanaceae. This understanding of the intraspecies structural variation can guide generation of mapping populations, and will facilitate the understanding of its genetic architecture for resistance to multiple diseases.

In this study, we also generated multi-omics datasets including resequencing data of 52 *S. americanum* accessions, RNAseq, SMRT-RenSeq and cDNA-RenSeq data. The resequencing data enabled us to understand the phylogeny and population structure of these accessions, and indicate that some accessions may belong to other species. Importantly, molecular markers can be rapidly developed by using the reference genomes and the resequencing data for any mapping populations. These data also enabled us to build a *S. americanum* Pan-NLRome to study the evolution and function of the *NLR* genes in *S. americanum*. Automatic annotation of NLR gene models remains a challenge, and therefore we manually annotated all *NLR* genes in the *S. americanum* reference genomes SP1102, SP2271 and SP2273. These gene models will facilitate curation of the *NLR* genes from other Solanaceae species, and will help the cloning and characterization of NLR immune receptors. Notably, consistent with previous findings ^40^, nearly half of the *S. americanum NLR* genes are in the *NRC* superclade. We found that an *NRC1* homolog that is lacking in the *N. benthamiana* genome, supports the function of Rpi-amr3 but not Rpi-amr1 (Supplementary Fig. 10), thus expanding our knowledge of the NRC network in Solanaceae. We also found that *NRC4b* clade has expanded in *S. americanum* compared to *N. benthamiana* and tomato ^41^. Whether these *NRC4* paralogs are functionally redundant or divergent requires further study.

Potato late blight triggered the Irish famine in the 1840s, and remains a global challenge that greatly constrains potato production. To understand effector-triggered immunity (ETI) of *S. americanum* to *P. infestans*, we used “effectoromics” to dissect the ETI interactions between them. Using transient expression of effectors, we generated a matrix of 315 RXLR effectors × 52 *S. americanum* accessions. As expected, most RXLR effectors cannot be recognized by most *S. americanum* accessions. Interestingly, AVRamr1 (36/52) and AVRamr3 (43/52) recognitions are widely distributed in *S. americanum* accessions, indicating that *Rpi-amr1* and *Rpi-amr3* play an important role in the late blight resistance of *S. americanum*. This finding is consistent with the conclusions of a Pan-genome ETI study of the interaction between *Arabidopsis* and effectors from diverse strains of *P. syringae* ^44^. Some effectors induce cell death in all *S. americanum* accessions, such as effectors in the AVRblb2 family (Fig. 3). This observation is consistent with previous findings that AVRblb2 (PexRD39/40) induces cell death in all tested wild potato species, but not in *N. benthamiana* ^45,46^. Conceivably, this non-specific cell death might be the result of the virulence activities of AVRblb2. Some resistant accessions lack AVRamr1 and AVRamr3 recognition, and thus are valuable sources of novel *Rpi* genes. However, isolation of novel *Rpi* genes from resistant accessions that carry *Rpi-amr1* and *Rpi-amr3* requires their removal by either genetic segregation or gene knockout.

In the current study, this challenge was successfully addressed using effectoromics. Using GWAS or BSA-RenSeq approaches, we cloned three novel *Rpi* genes: *Rpi-amr4, Rpi-amr16* and *Rpi-amr17*, as well as their cognate *Avr* genes *Avramr4* (*PITG_22825*), *Avramr16* (*PITG_02860*) and *Avramr17* (*PITG_04373*) (Fig. 4 and Fig. 5). Using the transient expression assay in *N. benthamiana*, we showed that *Rpi-amr4* contributes to *P. infestans* resistance. Using CRISPR-Cas9 in *S. americanum*, we generated *Rpi-amr4* and *Rpi-amr16* knockout SP2271 lines respectively, as well as the *Rpi-amr1*/*Rpi-amr3*/*Rpi-amr17* triple knockout SP2300 lines. These enabled us to validate gene function in *S. americanum* and create materials for cloning new *Rpi* genes. *Avramr4, Avramr16* and *Avramr17* are present in all tested *P. infestans* isolates, and are up-regulated upon infection ^31,47^. Unlike some RXLR effectors that belong to superfamilies such as *Avr2* and *Avrchc1*^12,48^, *Avramr4, Avramr16* and *Avramr17* are all singletons. AVRamr16 (PITG_02860) was reported to promote host susceptibility by targeting a host protein NLR1 ^42^, and the virulence function(s) and host targets of AVRamr4 and AVRamr17 remain to be discovered.

*Rpi-amr4-1102* confers quantitative and incomplete resistance in *N. benthamiana* transient assays compared to the full resistance conferred by *Rpi-amr3. Rpi-amr16* was isolated from SP2271-a late blight susceptible accession in detached leaf assays (DLAs). These newly isolated *Rpi* genes thus likely confer weaker late blight resistance compared to the previously cloned *Rpi-amr1* and *Rpi-amr3*, despite showing strong HR when transiently co-expressed with their cognate recognized effectors. However, many effectors are suppressors of host immunity, notably AVRcap1 that can attenuate the function of the helper NLRs NRC2 and NRC3 ^49^, and this may explain why some *Rpi* genes are weaker than others. Resistance based on recognition of a single effector can be easily overcome by mutations or silencing, as shown by *Avrvnt1* ^50,51^. *Rpi-amr4, Rpi-amr16* and *Rpi-amr17* belong to *Rpi-amr3, R3* and *Rpi-chc1* clades, respectively (Fig. 2a) and *R3* and *Rpi-chc1* are not NRC-dependent. Since resistance is a quantitative phenotype, as illustrated by the fact that most *R* genes are semi-dominant (resistance in heterozygotes is weaker than in homozygotes), and ETI functions at least in part by potentiating PTI ^52^, then more recognition capacity should strengthen resistance, even if some recognition capacities confer weak resistance alone. Even weak resistance genes can thus be useful, and the three newly cloned *Rpi* genes could be stacked with *Rpi-vnt1, Rpi-amr1* and *Rpi-amr3* to quantitatively increase defence activation ^52^, minimize the risk of *Rpi* gene breakdown, and ultimately help to turn potato into a nonhost plant of late blight as is *S. americanum*. We anticipate that additional effector-detecting immune receptor genes can be rapidly cloned using this pipeline.

Both Rpi-amr1 and Rpi-amr3 can recognize the AVRamr1 and AVRamr3 homologs from other *Phytophthora* pathogens, and *Rpi-amr3* confers resistance against *P. parasitica* and *P. palmivora* by detecting their conserved AVRamr3 homologs ^29,30^. Conceivably, Rpi-amr4, Rpi-amr16 and Rpi-amr17 might also recognize the effectors from other important *Phytophthora* pathogens and confer resistance against multiple diseases.

In summary, our study provides valuable genomic and genetics tools that accelerate the path to understanding and achieving durable resistance against potato late blight and other plant diseases, and make *S. americanum* an excellent model plant to study molecular plant-microbe interactions and plant immunity.

## Author contributions

X.L. Y.J., K.H.S, S.H. and J.D.G.J. conceived and designed the project. X.L. and Y.J. wrote the first draft with inputs from all the authors. X.L., Y.J., K.H.S., S.H. and J.D.G.J. reviewed and edited the manuscript. Y.J., M.P., X.L. and R.K.S. performed the bioinformatics analyses. X.L. and M.M., performed the effectoromics screening. X.L., R.H., M.S., A.C.O.A., S.F. and A.N. contributed to clone and characterize *Rpi-amr4, Rpi-amr16* and *Rpi-amr17*. M.Smoker and J.T. performed the plant transformation. K.W., Y.L., C.Z., S.J.P., K.H.S., S.H. and J.D.G.J. contributed resources.

## Acknowledgements

This research was financed from BBSRC grants BB/P021646/1, BB/S018832/1, BB/W017423/1, the Gatsby Charitable Foundation, National Key Research and Development Program of China (2019YFA0906200), Guangdong Major Project of Basic and Applied Basic Research (2021B0301030004), Agricultural Science and Technology Innovation Program (CAAS-ZDRW202101), National Research Foundation of Korea (2018R1A5A1023599 and 2019R1A2C2084705) and BioGreen21 Agri-Tech Innovation Program (PJ01579901). We thank TSL transformation team (Aleksandra Wawryk-Khamdavong), SynBio team (Mark Youles and Liam Egan), media services (Neil Stammars), bioinformatics team (Dan MacLean) and horticultural team (Sara Perkins, Justine Smith, Lesley Phillips, Catherine Taylor, Timothy Wells, Damian Alger and Sophie Able) for their support. We thank Sylvestre Marillonnet for sharing the Cas9 construct (pAGM47523). We thank Experimental Garden and Genebank of Radboud University, Nijmegen, The Netherlands, IPK Gatersleben, Germany and Sandra Knapp (Natural History Museum, London, UK) for access to *S. americanum* genetic diversity. We thank Vivianne G. A. A. Vleeshouwers at Wageningen University and Research, Paul Birch, Ingo Hein and Brian Harrower at James Hutton Institute for making available clones of some effectors. We thank Brande B.H. Wulff, Sanu Arora and Kumar Gaurav for helpful discussions.

## Competing interests

The authors declare no competing interests.

## Supplementary Information

**Supplementary Figure 1**. Phenotypes of four *S. americanum* accessions at the flowering stage.

**Supplementary Figure 2**. K-mer distribution of four *S. americanum* genomes.

**Supplementary Figure 3**. Hi-C interaction maps of SP1102, SP2271 and SP2273.

**Supplementary Figure 4**. BUSCO evaluation of *S. americanum* genome assemblies and gene model predictions.

**Supplementary Figure 5**. Genome alignment between *S. americanum* and neighbouring species. (a). Genome alignment between *S. americanum* (SP1102) and potato (DM). (b). Genome alignment between *S. americanum* (SP1102) and eggplant (HQ1315).

**Supplementary Figure 6**. Large structural variations between *S. americanum* genomes. Structural variations with length < 1 Mb were not shown in the figure.

**Supplementary Figure 7**. Structural variations among *S. americanum* genomes. (a). SV length distribution. (b). Number of SVs in each accession. (c). SV length in each accession.

**Supplementary Figure 8**. SVs might contribute to gene expression variations. (a). Number of SVs overlapped with genomic features. (b). An example of different expressed gene (DEG) whose expression might associate with the SV in its promoter region. Upper panel: The gene model of *sp1102chr11_nlr_6* with a deletion in its promoter region. Bottom panel: the volcano plot of DEGs in leaves between SP1102 and SP2271. Red and blue dots stand for DEGs (*P*-value < 0.05) with fold change <= 0.5 or >= 2, respectively; Green dots denote non-significantly expressed genes. The purple star: the expression of *sp1102chr11_nlr_6*.

**Supplementary Figure 9**. Sequence diversity in NLR region. (a). The comparison of sequence diversity between NLR regions and non-NLR regions. Wilcoxon rank-sum test was used to assess the diversity index between NLR regions and non-NLR regions. (b). Synteny plot of *Rpi-amr3* locus indicates the sequence diversity of NLR regions.

**Supplementary Figure 10**. SaNRC1 supports the function of *Rpi-amr3* but not *Rpi-amr1*; SaNRC2 supports the function of both Rpi-amr3 and Rpi-amr1. (a). HR assay in nrc2.3.4 knockout *N. benthamiana*; (b). HR assay in wild-type *N. benthamiana*. OD^_600_^ = 0.5.

**Supplementary Figure 11**. Venn plot for effector recognition profiles in the four *S. americanum* accessions with reference genomes.

**Supplementary Figure 12**. Phylogeny, population structure and inbreeding coefficient of all *S. americanum* accessions in this study.

**Supplementary Figure 13**. BSA-RenSeq for Rpi-amr4 in F2 population of SP2271 x SP2298. (a). Pipeline of the BSA-RenSeq; (b and c) Informative SNPs from the BSA-RenSeq are located on a NLR singleton sp1102chr01_nlr39.

**Supplementary Figure 14**. Alignment of Rpi-amr4-1102 and Rpi-amr4-2271 proteins.

**Supplementary Figure 15**. PITG_22825 agroinfiltration and disease test on the *Rpi-amr4* transgenic *N. benthamiana*.

**Supplementary Figure 16**. *Rpi-amr16* knockout SP2271 lines. (a). The constructs used for the CRISPR-Cas9 knockout experiment; (b). Genotyping data of three *Rpi-amr16* lines SLJ25598#1, SLJ25598#2 and SLJ25598#3. (c). Phenotype of three selected knockout lines by agroinfiltration. Rpi-amr3 and AVRamr3 were used as controls.

**Supplementary Figure 17**. *Rpi-amr1*/*Rpi-amr3*/*Rpi-amr17* triple knockout SP2300 lines. (a). The constructs used for the CRISPR-Cas9 knockout experiment; (b, c and d). Genotyping the *Rpi-amr1-2300* and *Rpi-amr3-2300* and *Rpi-amr17* in two knockout lines SLJ25603#3 and SLJ25603#17. (e). Phenotype of the triple knockout lines after expression of PpAVRamr1, AVRamr3 and PITG_04373. Co-expression of Rpi-amr17 and PITG_04373 was used as a control. (f). Disease test on the two triple knockout lines, *P. infestans* isolates T30-4 and 88069 was used in this assay. SP2300 and SP2271 were used as controls.

**Supplementary Dataset 1**. ORFs of the manually annotated *NLRs* from SP1102.

**Supplementary Dataset 2**. CDSs of the manually annotated *NLRs* from SP1102.

**Supplementary Dataset 3**. Proteins of the manually annotated *NLRs* from SP1102.

**Supplementary Table 1**. NLR physical position, P/A polymorphism and cDNA RenSeq data.

**Supplementary Table 2**. Effectoromics screening scoring table.

**Supplementary Table 3**. Primers and molecular markers in this study.

**Supplementary Table 4**. Genotyping data of *Rpi-amr4* knockout SP2271 lines.

## Notes

### Competing Interest Statement

The authors have declared no competing interest.

