## Supplemental Figure 1-17 for "The *Solanum americanum* pangenome and effectoromics reveal new resistance genes against potato late blight"

### Supplemental Information

#### Supplemental Methods

##### Sequencing and assembly *S. americanum* genomes

Four representative *S. americanum* accessions, SP1102, SP2271, SP2273 and SP2275 were selected for sequencing. The Pacific Bioscience Sequel II platform in the circular consensus sequencing (CCS) mode was applied to sequence the genomes of SP1102 and SP2271 and generated 30.5 Gb and 28.5 Gb high-fidelity (HiFi) reads, respectively. The PromethION and GridION platforms of Oxford Nanopore Technologies were applied to sequence the genomes of SP2273 and SP2275, and generated ~81.1 Gb and ~114.5 Gb data, respectively. To estimate the genome heterozygosity and polish the raw assembled genomes, we also prepared the libraries for Illumina pair-end short reads sequencing following the standard protocol and generated an average of 99.2 Gb clean data for each Sam accession by Illumina Hiseq 2500 platform. The Hi-C libraries of three Sam accessions, SP1102, SP2271 and SP2273, were created from young seedlings based on the restricted enzyme MboI. Illumina Hiseq 2500 platform was applied to generate 86.5, 81.8 and 54.8 Gb pair-end reads for SP1102, SP2271 and SP2273, respectively.

The genome size and heterozygosity were estimated using a k-mer-based approach by KAT [1] and GenomeScope [2]. The Hifiasm [3] was applied to *de novo* assemble SP1102 and SP2271 using default parameters. Pseudo-chromosomes were built by using the juicer [4] and 3d-DNA[5] pipeline with parameters "-m haploid -i 15000 -r 0". To assemble the genome of SP2273 and SP2275, we first corrected ONT reads using Canu [6] with parameters 'correct corOutCoverage=500 corMinCoverage=2 minReadLength=2000 genomeSize=1g -nanopore-raw'. The corrected reads were then assembled into raw contigs by SMARTdenovo [7] with the following command line arguments 'perl smartdenovo.pl -c 1 -t 24 -k 17'. The raw assemblies were then iteratively polished using Illumina short reads. Reads were aligned to the raw assemblies using BWA [8] and resulting bam files were passed to Pilon [9] for polishing. The quality of assemblies was assessed by BUSCO (Benchmarking Universal Single-Copy Orthologs) [10] with solanales_odb10 database.

**Protein-coding genes prediction**

For SP1102 and SP2271, to help with gene model prediction, the transcriptomes of *S. americanum* whole seedlings, roots, stems, leaves, flowers and fruits were sequenced by using Illumina Hiseq 2500 platform with three replications for each tissue and 4 Gb clean data for each sample. The reads were aligned to the genome by HISAT [11], transcripts were assembled by StringTie [12], Cufflinks [13] and Trinity [14] respectively, then imported into PASA [15] for protein-coding gene prediction. *Ab initio* and homologous protein search strategies were also performed by using SNAP [16], AUGUSTUS [17], GlimmerHMM [18] and exonerate [19]. All predicted evidence was integrated by EVM [15]. To predict gene models in SP2273 and SP2275, we used ITAG4.0 [20] and SolTub_3.0 [21] datasets for homology-based gene prediction by GeMoMa program [22]. RNA-seq data, obtained from SP2273 was also incorporated for splice site prediction.

**Phylogenetic analysis of *S. americanum***

The representative protein sequences of *A. thaliana*, *P. inflata*, *C. annuum*, *S. melongena*, *S. tuberosum* Group Phureja (DM1-3 516 R44), *S. tuberosum* Group Tuberosum (RH89-039-16), *S. commersonii*, *S. chacoense*, *S. pennellii*, *S. pimpinellifolium*, *S. lycopersicum* and four *S. americanum* (SP1102, SP2271, SP2273 and SP2275) were extracted and input into OrthoFinder [23] to cluster orthogroups with MCL algorithm. The protein sequences of 1,363 single-copy orthogroups were extracted to infer the phylogenetic relationship following the supermatrix method.

Sequences from 15 genomes were aligned by MAFFT [24] with parameter '--auto' and trimmed by trimAl with '-phylip -gt 0.8'. IQ-TREE [25] was applied to infer the phylogenetic relationships with parameters '--alrt 1000 -B 1000'. We used BASEML and MCMCTREE from PAML package [26] to estimate the divergence time. The CDS sequences of 1,363 single-copy orthogroups were extracted for rough estimation of the substitution rate using BASEML with model = 7. MCMCTREE with parameters 'model = 7, burnin = 5,000,000, sampfreq = 300, nsample = 20,000' was applied to estimate the divergence time. The divergence time of potato - tomato (7 - 10 MYA) [27] and potato - Arabidopsis (111 - 131 MYA, http://www.timetree.org/) was used for calibration. Two rounds of estimation were performed with very similar results.

**Genomic alignments of Sam and neighboring species**

The genomic alignments between SP1102 and eggplant/potato were performed and by MUMMER [28] with parameters ' --maxmatch -c 100 -b 500 -l 50 '. The alignment was further filtered with ' -1 -i 80 -l 100' parameters. The delta format was then converted to PAF format by paftools.js script [29] and passed to D-Genies[31] for dot plot visualization. The same pipeline was used for genomic alignments among Sam genomes with modified parameters '--batch 1 -t 40 -l 100 -c 500 ' and '-1 -i 90 -l 100'.

**Syntenic analysis of Sam and neighboring species**

We applied MCscan [30] in Python version to perform the syntenic analysis. The representative protein sequences and corresponding gene model annotations in BED format of potato (DMv6.1), eggplant (HQ-1315) and four Sam genomes were extracted for searching homologous with '-m jcvi.compara.catalog ortholog --cscore=.99' parameters. The syntenic regions were identified with '-m jcvi.compara.synteny screen --minspan=30' and visualized with '-m jcvi.graphics.karyotype' parameters.

**Identification of structural variations**

To identify large structural variations (SVs, > 1Mb in length), we aligned the chromosome-grade assembly (SP2271 and SP2273) to the reference genome SP1102 by using MUMMER with parameters: '--batch 1 -t 20 -l 100 -c 500' and further filtered the alignment with parameters: '-i 90 -l 100'. SyRI v1.4 was adopted to identify SVs based on the alignment delta files, only large SVs were kept for further analysis.

The assembly-based approach was applied to identify structural variations (> 40 bp in length) among Sam genomes following the pipeline of SVIM-asm [31]. The contig assemblies of SP2271, SP2273 and SP2275 were aligned to SP1102 reference using minimap2 [29] with the following parameters '--paf-no-hit -a -x asm5 --cs -r2k'. SVs, which consist of insertions, deletions, duplications and inversions were identified using SVIM-asm with 'haploid' mode. The SVs were further annotated by SnpEff [32].

**Calculation of variation index across *S. americanum* genome**

To describe the variations across *S. americanum* genome, we introduced the variation index. We used the SV information generated by SVIM-asm and a slide-window (window = 500 kb, step = 50 kb) method for index calculation. The index of a window = (sum of SV length in a window) / window length. The final index of each window was generated from the average of SP2271, SP2273 and SP2275. Higher index value referring to higher variation level of a window. If a window overlapped an NLR gene, this window was counted as NLR-region, a total of 2,304 NLR-regions were extracted from the SP1102 genome. To compare the variations between NLR-region and non-NLR-region, we randomly selected 2,304 non-NLR-regions and compared variation index with NLR-regions by Wilcoxon rank-sum test. Ten rounds of randomly selection and comparison were performed.

**Annotation of NLR gene models**

The *NLR* genes from the 4 Sam reference genomes were predicted by NLR-annotator [33]. To get a better gene model for these *NLR* genes, all the *NLR* genes from SP2271, SP1102 and SP2273 genomes were manually curated. In brief, the outputs of NLR-annotator were imported into Geneious (v10.2.6) [34] as annotations of the reference genomes, the predicted *NLR* fragments with 2 kb flanking sequences from both sides were extracted, then Augustus[17] was used to predict the gene model based on the trained dataset of tomato. The gene models were curated based on functionally validated *NLR* genes from public databases, and cDNA RenSeq data were also used to assist the manual annotation.

**Phylogenetic analysis of NLRs in Sam genomes**

To infer the phylogeny of NLRs, the protein sequences of NB-ARC domain from NLR-annotator were aligned by MAFFT [24], and IQ-TREE was used to build the phylogenetic tree, JTT+F+R9 model was selected by ModelFinder [35], and used for inferencing the maximum likelihood tree, ultrafast bootstrap (UFBoot) [36] was set to 1000. The *CED-4* from *C. elegans* was selected as an outgroup.

To analyze the NLR presence and absence in Sam genomes, we collected 12 previously reported SMRT RenSeq assemblies[37] and generated three new assemblies from accessions SP2298, SP3370, SP2308. We used GMAP [38] to predict the NLR homologs among Sam genomes and SMRT RenSeq assemblies. The CDS sequences of manually curated NLRs in SP1102 genome were extracted and mapped to the rest 3 *S. americanum* genomes and 16 SMRT-RenSeq assemblies using GMAP with parameters '-f 2 -n 1 --min-trimmed-coverage=0.70 --min-identity=0.70'. The NLRs which failed to be aligned were marked as absent. In the v4 RenSeq library[39], more baits were included compared with the v3 RenSeq library[40], thus, if a certain NLR was absent in all v3 RenSeq assemblies but present v4 assemblies, the absence might be false-positive and marked as NA. To calculate the expression level of *NLRs* of SP1102, the RNA from young leaves of SP1102 were isolated and cDNA RenSeq was performed as described previously [41]. We mapped the reads of SP1102 cDNA RenSeq to its genome using STAR (2.6.0c) [42], the BAM files were imported into Geneious (v10.2.6) [34] and the TPM (Transcripts Per Kilobase Million) of *NLR* genes were calculated by the "Calculate Expression Levels" function. The NLR phylogeny, TPM and PAV results were passed to the online software iTOL[43] for final visualization.

**Re-sequencing of 52 Sam accessions**

The genomic DNA of 4-week-old young leaves from 52 Sam accessions were sampled and isolated by Qiagen DNeasy plant kit (Qiagen, 69104). Whole-genome PCR-free, 2 x 150 bp paired-end illumina library were generated and sequenced by Novogene (Beijing, China), and generated ~10 GB data for each Sam accession. The raw reads were trimmed by trimmomatic v0.36 [44]. The clean reads of each accession were mapped to the reference genome of SP1102 by minimap2 (v2.16) [29], and converted to BAM format by samtools (v1.9). The SNP calling was carried out by samtools and bcftools (v1.9).

To infer the phylogenetic relationships of Sam accessions, we selected the genomes of potato (DM 1-3 516 R44 v6.1), tomato (Heinz 1706 v4.0) and eggplant (v3) as an outgroup. Wgsim (https://github.com/lh3/wgsim) was used to simulate the whole-genome sequencing reads from potato, tomato and eggplant genomes with parameters: '-e 0 -d 350 -N 500000000 -1 150 -2 150 -r 0 -R 0 -X 0'. The simulated reads mapping and SNP calling were performed using the same approaches. Bedtools (v2.17) was used to extract SNPs in coding region. The SNP-based phylogenetic tree was inferred by IQ-TREE with 1000 ultrafast bootstrap and TVMe+R2 model, which was automatically selected by ModelFinder. The phylogenetic tree was visualized by FigTree (v1.4.4).

**GWAS analysis**

For the GWAS analysis, all the SNPs resided in the *NLR* gene region, as well as 3 kb upstream and 1kb downstream region, were extracted by bedtools (v2.17). The SNPs was filtered and processed by Plink (v1.90) with parameters '--make-bed --allow-extra-chr --allow-no-sex --mind 1 --maf 0.05 -geno 0.05 --recode --out'. The responsiveness scores of each effector were used as the phenotype and passed to Plink for association analysis with parameters '--allow-extra-chr --allow-no-sex –assoc --bfile --pheno'. The Manhattan plot was visualized by an R package qqman (v0.1.8).

**Effectoromics screening**

An RXLR effector library of 311 RXLR effectors was used in the effectoromics screening. The signal peptides were removed, and the effector domains were cloned into overexpression vectors (pMDC32 or pICSL86977) or PVX vectors. The *S. americanum* plants were grown in a containment glasshouse. Four or five-week plants were used for the agroinfiltration. For the overexpression vectors, the cell death was scored at 4 days after infiltration (dpi); for the PVX vectors, the cell death was scored at 7 days after infiltration (dpi). $OD_{600}$=0.5. The cell death phenotype was scored (Strong HR=2; Weak HR= 1; no HR=0). Two-leaves/plant, and two plants were used for each experiment.

**Constructs for transient overexpression**

To verify the candidate genes, the ORF of the candidate genes was amplified by Phusion high-fidelity DNA polymerase (NEB, M0530S) or KAPA HiFi Uracil+ DNA polymerase (Roche, 07959052001), then cloned into an overexpression vector pICLS86922 with 35S promoter and Ocs terminator by *BsaI* (NEB #R3733) or a USER cloning vector pICSLUS0004OD with 35S promoter and Ocs terminator by USER enzyme (NEB #M5508). The verified constructs were transformed into *Agrobacterium* for transient expression *in planta*.

#### Gene knockout by CRISPR-Cas9 system

For the knockout constructs, the guild RNAs were designed in Geneious (v10.2.6) by the "Find CRISPR Site" function with parameters: 'Maximum mismatches allowed against off-targets = 3; Maximum mismatches allowed to be indels = 0; pair CRISPR Sites: Maximum overlap of paired sites = 100; Maximum allowed space between paired sites = 300'. The reference genome or SMRT-RenSeq assembly was used for scoring against the off-targets activity. The selected guild RNAs are shown in Table S3. Two guild RNAs for each candidate gene were amplified with the sgRNA scaffold by Q5 high-fidelity DNA polymerase (NEB, M0491S), and vector pICSL70001 was used as the template. The fragments were then fused with an Arabidopsis U6-26 promoter (pICL90002) and cloned into Level 1 vectors of different positions (Position 3: pICH47751, Position 4: pICH47761, Position 5: pICH47772, Position 6: pICH47772). For the final level 2 constructs, *Cas9* with intron (position 1: pICSL11197), NPTII (position 2: pICSL11055), an end linker pICH41922, and the guild RNAs were assembled into pICSL4723_OD. The final constructs were then transformed into *S. americanum* accessions from gene knockout. After transformation, the T0 lines were moved into a containment glasshouse for phenotyping and genotyping. Agroinfiltration of the corresponding effector was used for the phenotyping. The gDNA from the individual T0 lines were isolated, and specific primers were designed for the *Cas9* gene and the target genes. The amplicons from the target genes were sub-cloned into a TA clone vector pGEM-Teasy (Promega, A1360) or pICSL86977 for sequencing. The sequencing data was analyzed in Geneious (v10.2.6).

#### Plant growth and transformation

The *N. benthamiana* and *N. tabacum* cv. Petit Gerard plants were sowed and grown in a controlled environment room (CER) with 22 °C, 45-65% humidity, and 16 hours of photoperiod. Four-week plants were used for the HR assay.

For the *S. americanum* transformation, the sterilized seeds (SP2271 and SP2300) were sown in MS medium (2% sucrose). The leaf discs were cut from 4-6 weeks old in-vitro plants. Add 100 µl of overnight *Agrobacterium tumefaciens* (AGL1) culture and 200 µM Acetosyringone into 20 ml of LSR broth, gently dip the leaf discs into the solution using sterile forceps for 20 minutes. Then remove the leaf discs from the *Agrobacterium tumefaciens* suspension, blot dry, and incubate under low light conditions at 18-24 °C for 3 days. Plate them on LSR1 + 200 µM Acetosyringone solid media. Transfer co-cultivated explants to LSR1 medium with selection antibiotics in the petri dish (about 7 leaf discs per plate). Subculture the explants onto fresh LSR1 media approximately every 14 days. Once the calli have sufficiently developed transfer them onto LSR2 media. Keep subculturing the explants every 14 days and shoots will start to appear. Remove shoots with a sharp scalpel and plant into MS2R solid media with selection antibiotics. Transgenic plants harboring appropriate antibiotic or herbicide resistance genes should have rooted normally by the fourth week and can be weaned out of tissue culture into sterile peat blocks before being transplanted to the glasshouse. LSR broth (1x Murashige and Skoog medium, 3% Sucrose, pH 5.7); LSR1 medium (1x Murashige and Skoog medium, 3% Sucrose, 2.0 mg/L zeatin riboside, 0.2 mg/L NAA, 0.02 mg/L GA3, 0.6% Agarose, pH 5.7); LSR2 medium (1x Murashige and Skoog medium, 3% Sucrose, 2.0 mg/L zeatin riboside, 0.02 mg/L GA3, 0.6% Agarose, pH 5.7); MS2R (1x Murashige and Skoog medium, 2% Sucrose, 100 mg/L Myo-inositol, 2.0 mg/L glycine, 0.2% Gelrite, pH 5.7).

#### Disease assay

*Phytophthora infestans* isolates T30-4, 88069 and NR9 4HH were used for the disease test, they were maintained on rye sucrose agar (RSA) medium in an 18°C incubator. To induce zoospores, ice-cold water was added to the 10-14 days-old plates. The plate was then incubated at 4 °C for 1-2 hours, then a hemocytometer was used to measure the number of zoospores. The zoospore suspension was used for the detached leave assay (100-500 zoospores/droplet).

#### BSA-RenSeq and map-based cloning

Three mapping populations were used in this study, they are F2 population of SP2271 x SP2272; BC1 and F2 populations of SP2271 x SP2300. The populations were phenotyped by agroinfiltration of RXLR effectors. A cork borer was used for sampling, and the leave disks from the responsive and non-responsive progenies were pooled respectively. The gDNA was isolated by Qiagen DNeasy plant kit (Qiagen, 69104). The RenSeq libraries were then prepared, as described previously [45]. The libraries were sequenced (Illumina 2 x 250 bp reads) in Novogene (Beijing, China). The SNP filtering and calling steps were described previously [46].

To design molecular markers, the 10x PCR-free re-sequencing reads were mapped to the Sam reference genome SP2271. Then SCAR markers that linked with the BSA-RenSeq signals were designed, the amplicons should only be present in the non-responsive allele. The SCAR markers were firstly tested on the parental lines, the verified markers were then used on gDNA from individual non-responsive plants. GoTaq G2 DNA polymerase (Promega, 0000066542) was used for the genotyping.

#### Data availability

All of the raw sequencing data generated in this study have been deposited at the National Center for Biotechnology Information (NCBI) Sequence Read Archive (SRA) with BioProject accession number PRJNA845062 (https://dataview.ncbi.nlm.nih.gov/object/PRJNA845062?reviewer=hliiufd2hm679172evsbdgcr69). The assembled genomes and annotations are available at Figshare (https://figshare.com/articles/dataset/The_Solanum_americanum_pangenome_and_effectoromics_reveal_new_resistance_genes_against_potato_late_blight/20454432).

**Supplemental Figures:**

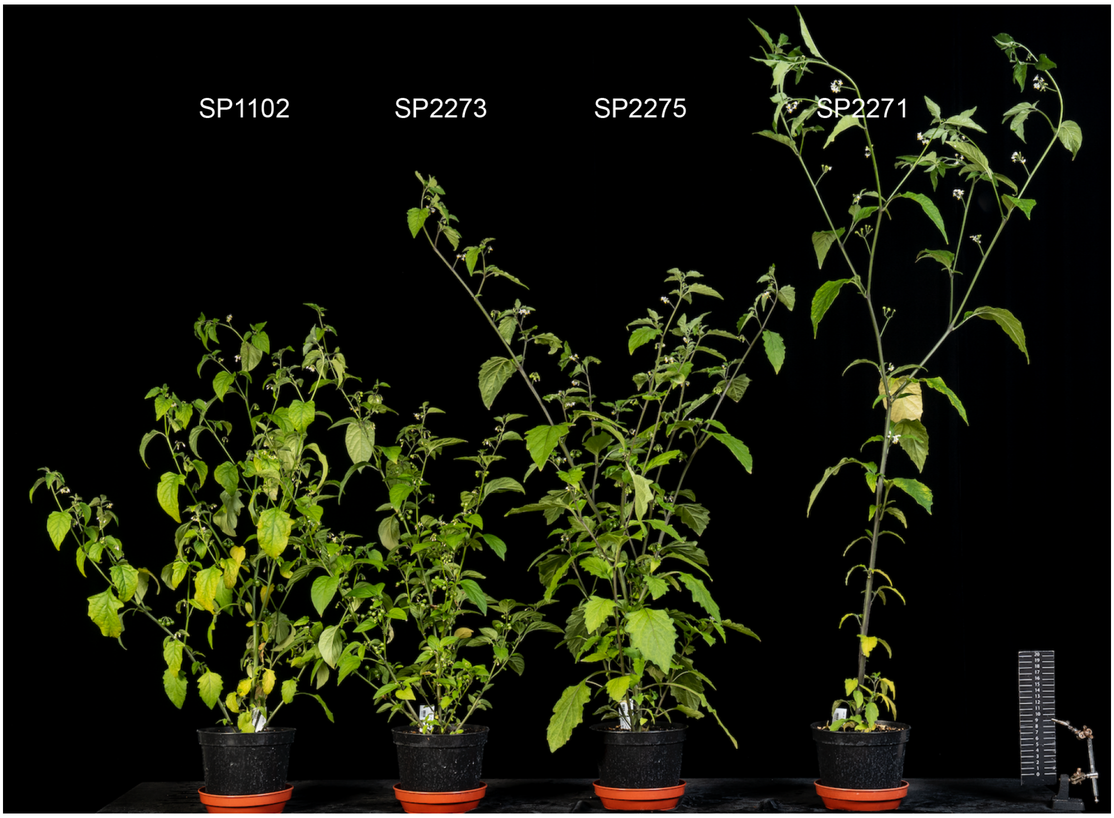


**Supplementary Figure 1.** Phenotypes of four *S. americanum* accessions at the flowering stage.

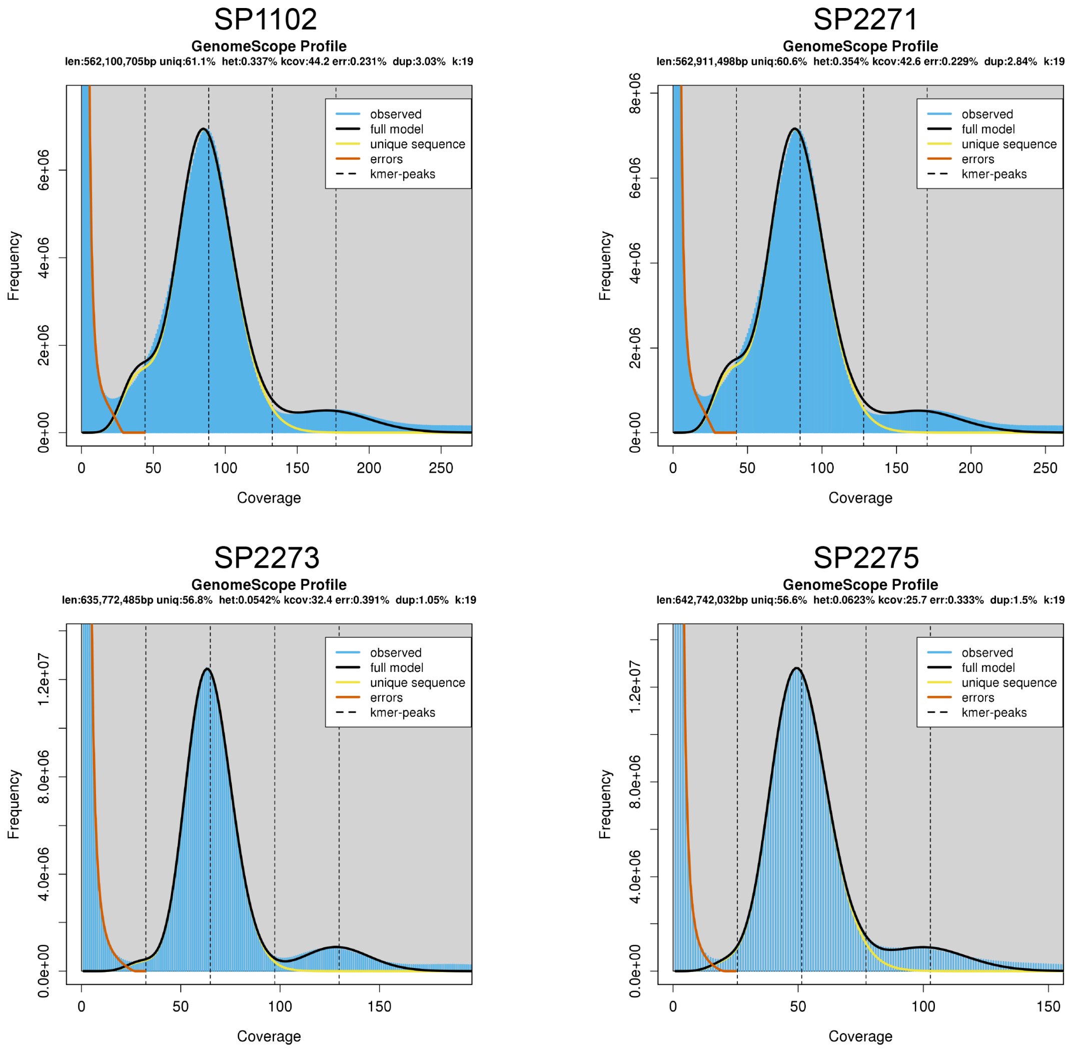


**Supplementary Figure 2**. K-mer distribution of four *S. americanum* genomes.

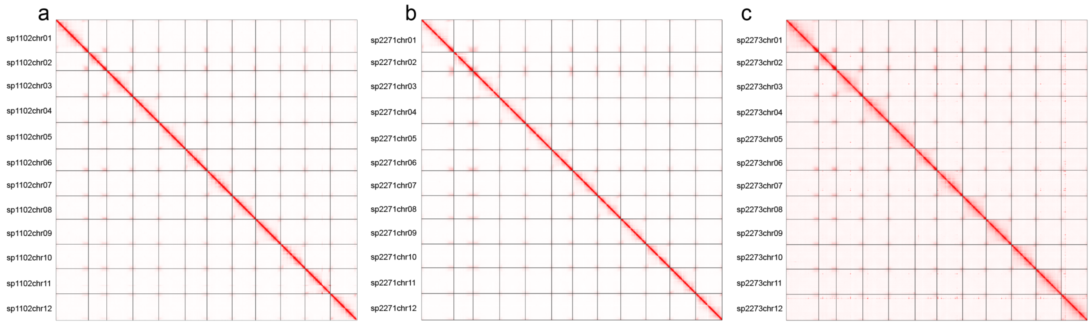


**Supplementary Figure 3**. Hi-C interaction maps of SP1102, SP2271 and SP2273.

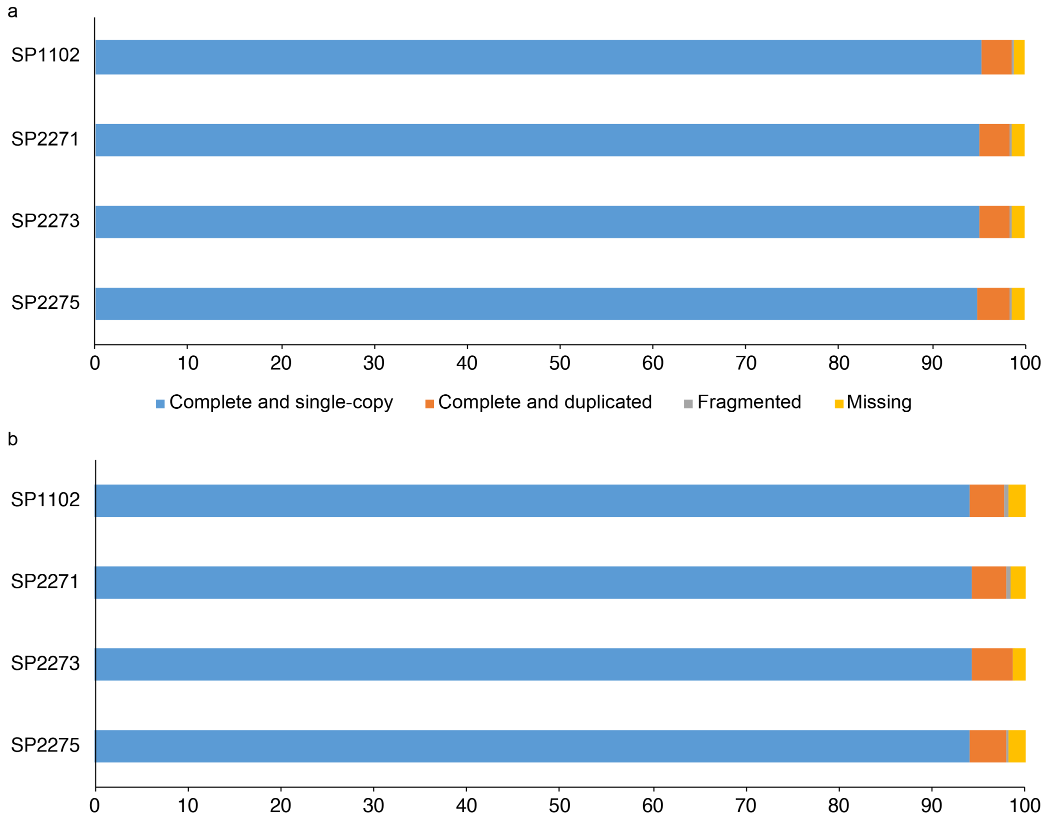


**Supplementary Figure 4**. BUSCO evaluation of *S. americanum* genome assemblies and gene model predictions.

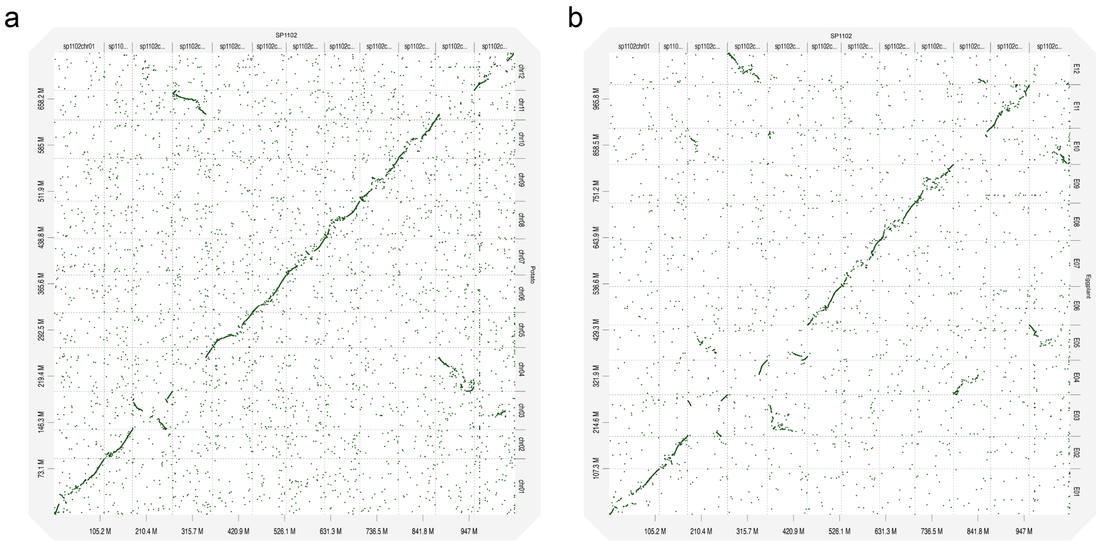


**Supplementary Figure 5**. Genome alignment between *S. americanum* and neighbouring species. (a). Genome alignment between *S. americanum* (SP1102) and potato (DM). (b). Genome alignment between *S. americanum* (SP1102) and eggplant (HQ1315).

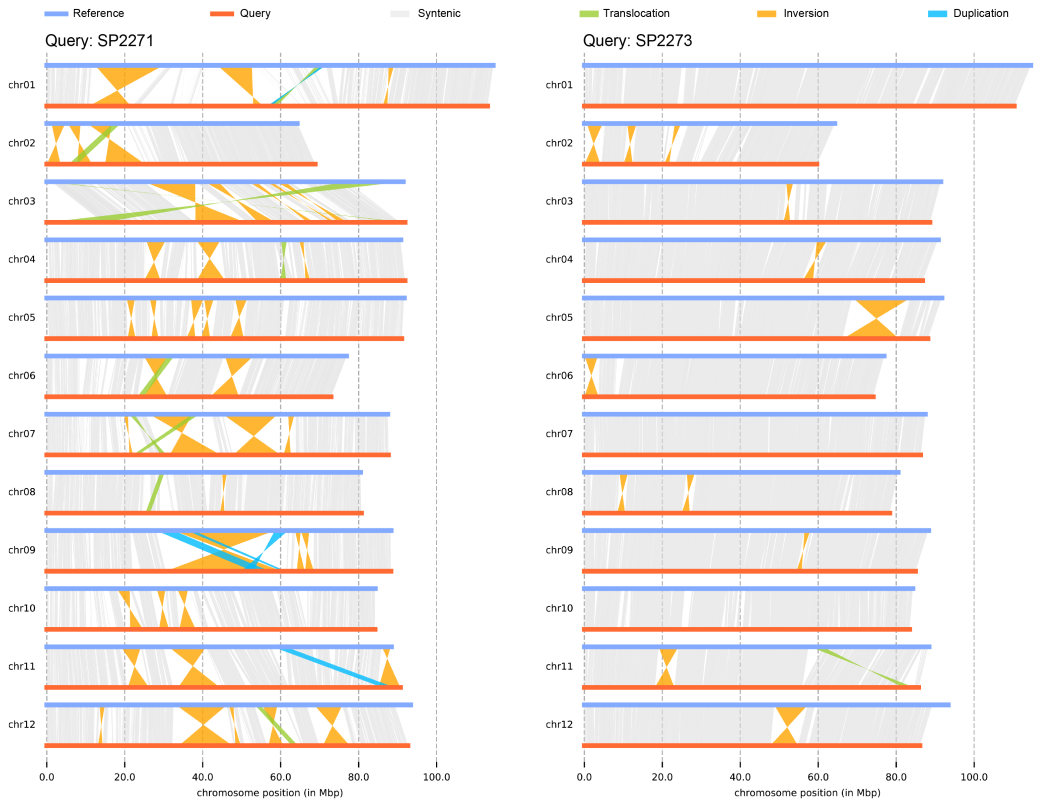


**Supplementary Figure 6**. Large structural variations between *S. americanum* genomes. Structural variations with length < 1 Mb were not shown in the figure.

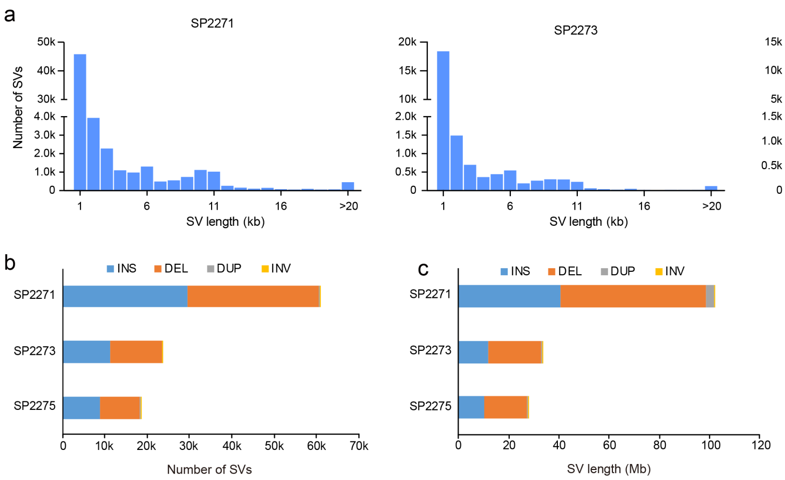


**Supplementary Figure 7**. Structural variations among *S. americanum* genomes. (a). SV length distribution. (b). Number of SVs in each accession. (c). SV length in each accession.

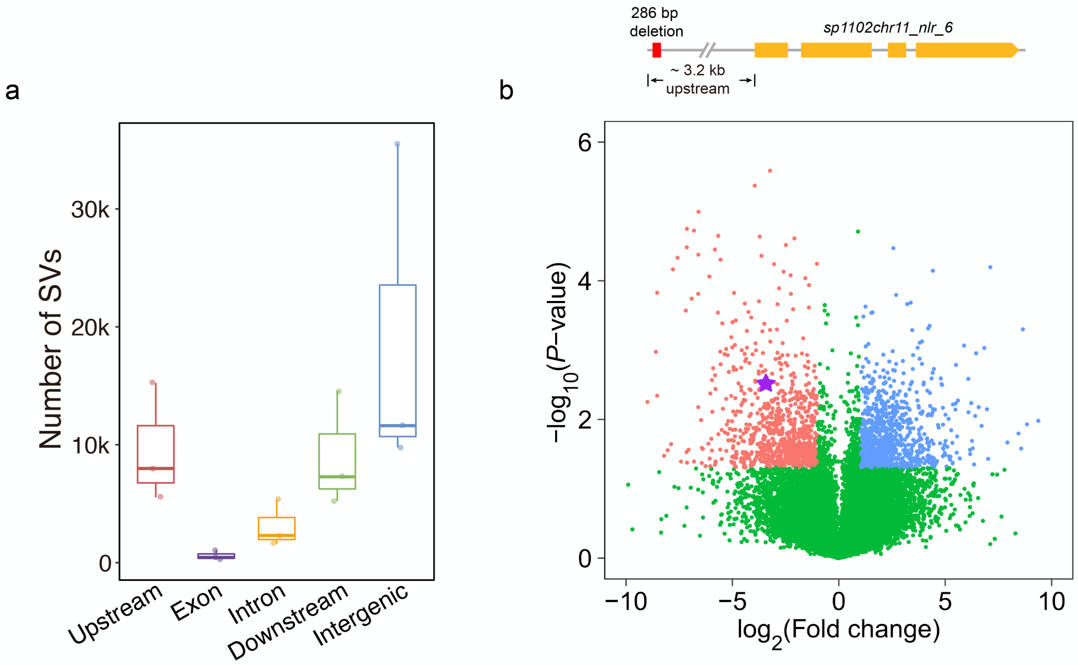


**Supplementary Figure 8**. SVs might contribute to gene expression variations. (a). Number of SVs overlapped with genomic features. (b). An example of different expressed gene (DEG) whose expression might associate with the SV in its promoter region. Upper panel: The gene model of *sp1102chr11_nlr_6* with a deletion in its promoter region. Bottom panel: the volcano plot of DEGs in leaves between SP1102 and SP2271. Red and blue dots stand for DEGs (*P*-value < 0.05) with fold change <= 0.5 or >= 2, respectively; Green dots denote non-significantly expressed genes. The purple star: the expression of *sp1102chr11_nlr_6*.

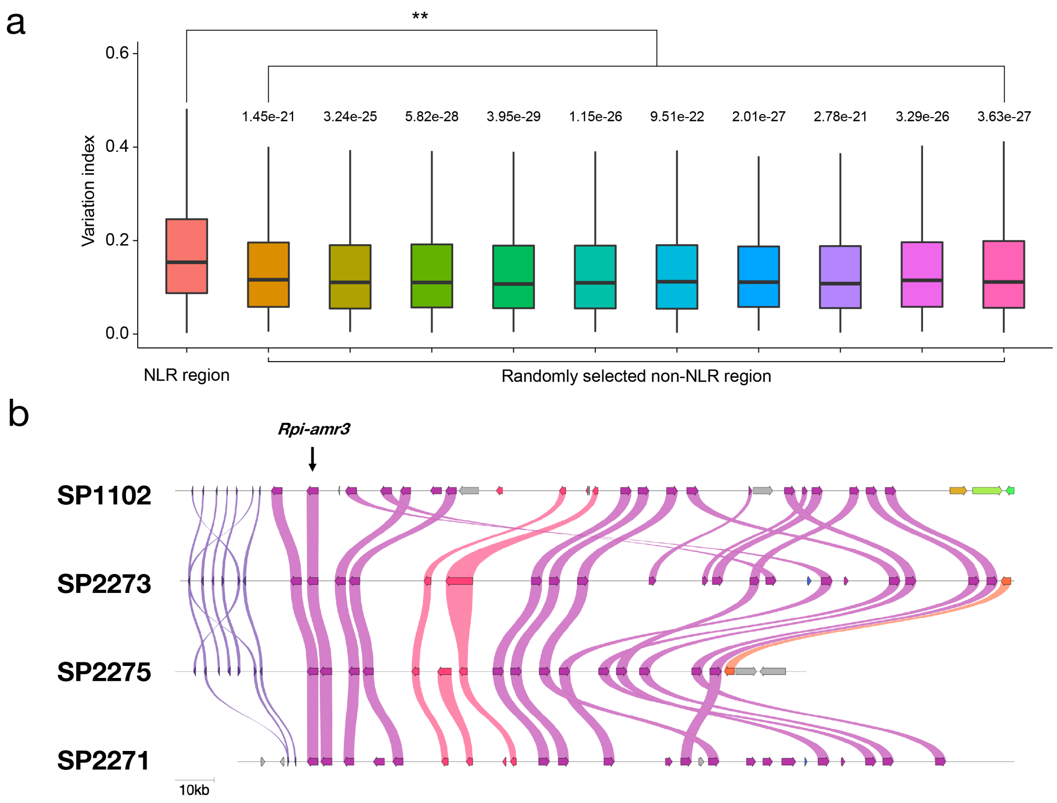


**Supplementary Figure 9**. Sequence diversity in NLR region. (a). The comparison of sequence diversity between NLR regions and non-NLR regions. Wilcoxon rank-sum test was used to assess the diversity index between NLR regions and non-NLR regions. (b). Synteny plot of *Rpi-amr3* locus indicates the sequence diversity of NLR regions.

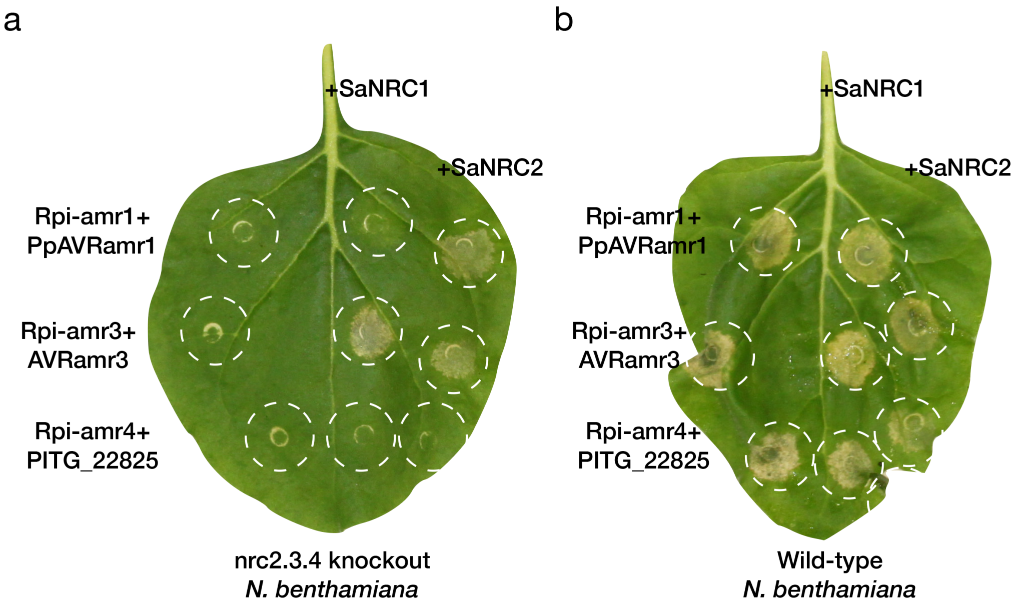


**Supplementary Figure 10**. SaNRC1 supports the function of *Rpi-amr3* but not *Rpi-amr1*; SaNRC2 supports the function of both Rpi-amr3 and Rpi-amr1. (a). HR assay in nrc2.3.4 knockout *N. benthamiana*; (b). HR assay in wild-type *N. benthamiana.* $OD_{600}$ = 0.5.

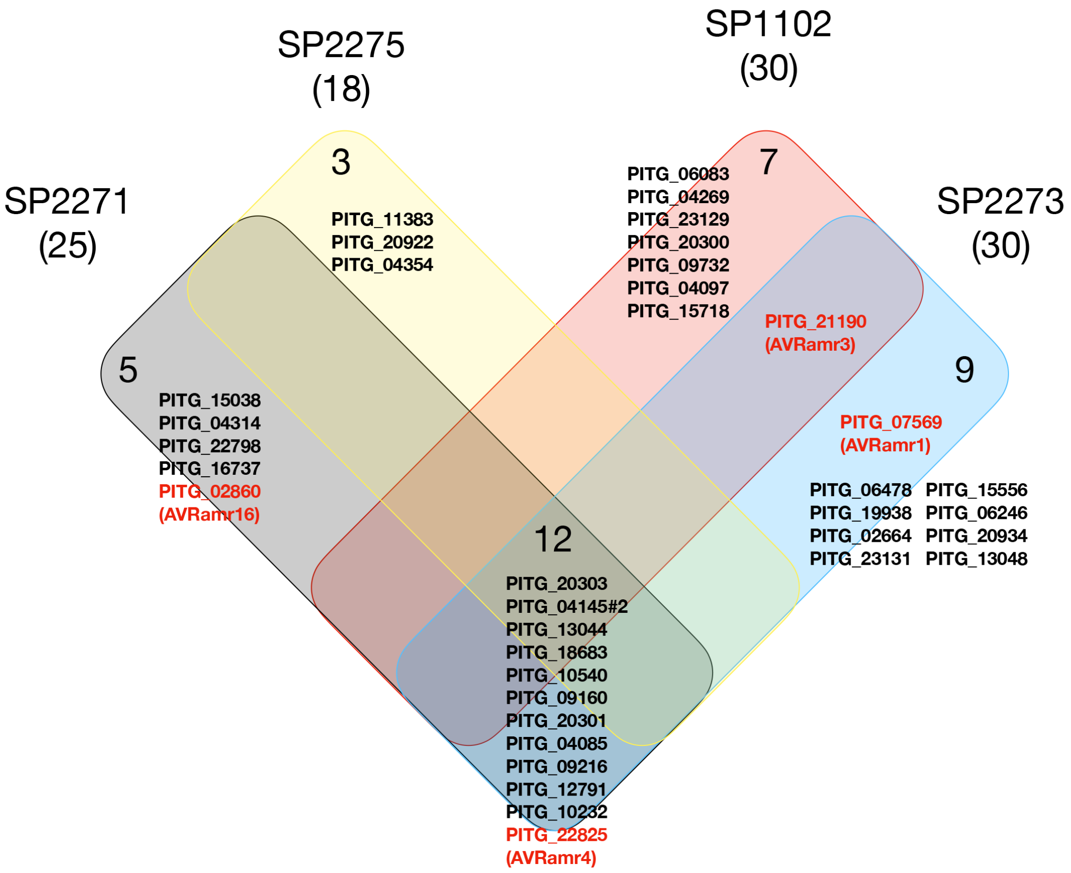


**Supplementary Figure 11**. Venn plot for effector recognition profiles in the four *S. americanum* accessions with reference genomes.

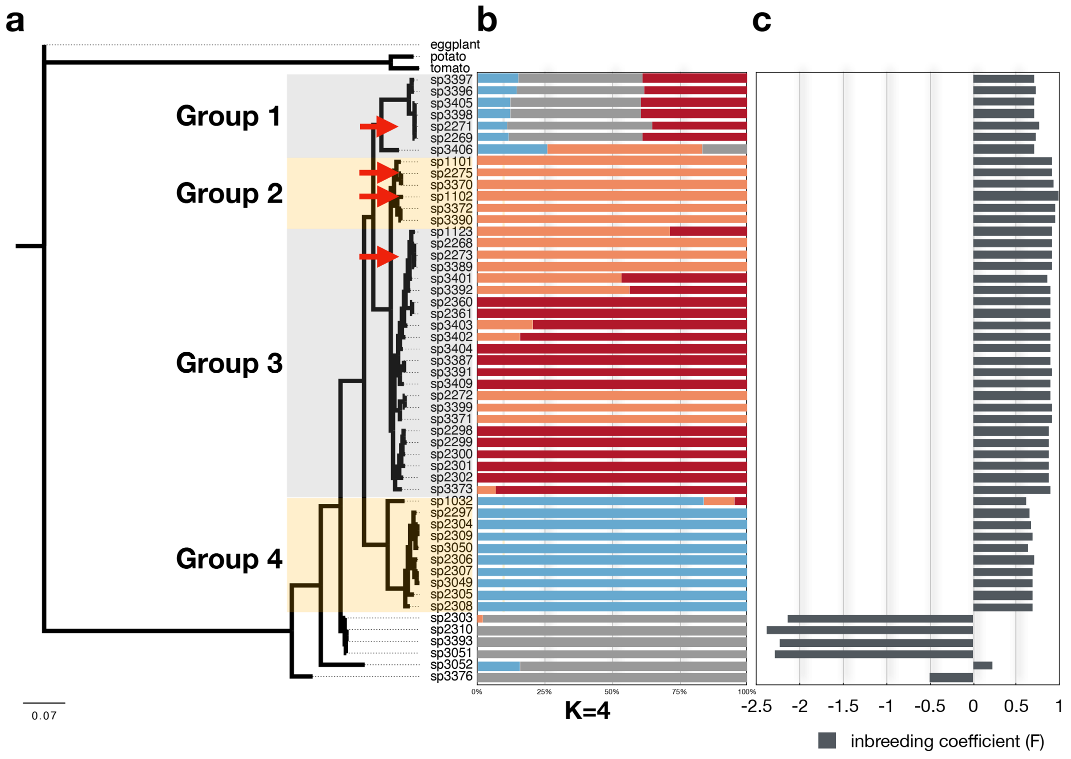


**Supplementary Figure 12**. Phylogeny, population structure and inbreeding coefficient of all *S. americanum* accessions in this study.

a

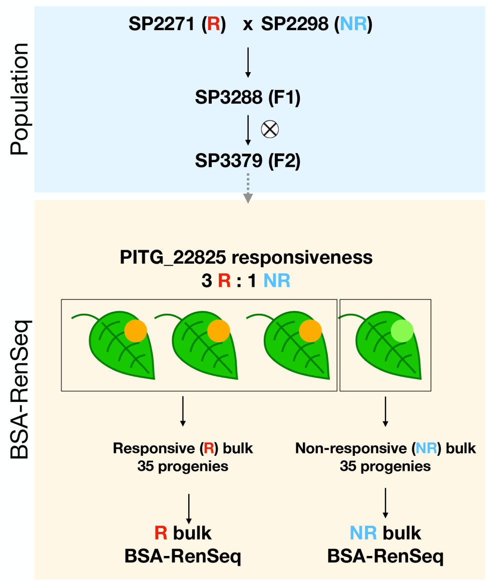


b

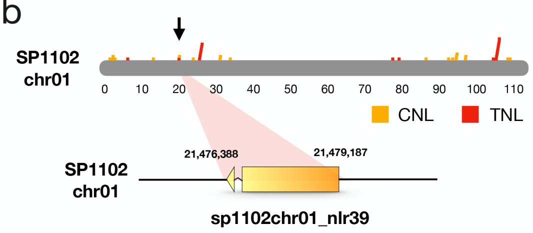


c

| Chromosome | Position | REF | ALT | FREQ_NR | FREQ_R |
|---|---|---|---|---|---|
| sp1102chr01 | 7451060 | C | T | 98.33 | 23.62 |
| sp1102chr01 | 21474598 | G | A | 98.68 | 32 |
| sp1102chr01 | 21474835 | T | G | 100 | 33.33 |
| sp1102chr01 | 21475956 | T | C | 100 | 37.39 |
| sp1102chr01 | 21477225 | G | A | 100 | 38.98 |
| sp1102chr01 | 21477436 | A | C | 100 | 31.94 |
| sp1102chr01 | 21477450 | A | C | 100 | 32.88 |
| sp1102chr01 | 21477592 | G | C | 95.59 | 24.3 |
| sp1102chr01 | 21477608 | T | G | 98.48 | 25.93 |
| sp1102chr01 | 21477703 | T | A | 100 | 24.39 |
| sp1102chr01 | 21477754 | G | C | 98.61 | 28.57 |
| sp1102chr01 | 21478376 | A | C | 100 | 41.94 |
| sp1102chr01 | 21478721 | G | A | 100 | 39.34 |
| sp1102chr01 | 78572703 | G | C | 98.7 | 36.51 |

**Supplementary Figure 13**. BSA-RenSeq for Rpi-amr4 in F2 population of SP2271 x SP2298. (a). Pipeline of the BSA-RenSeq; (b and c) Informative SNPs from the BSA-RenSeq are located on a NLR singleton sp1102chr01_nlr39.

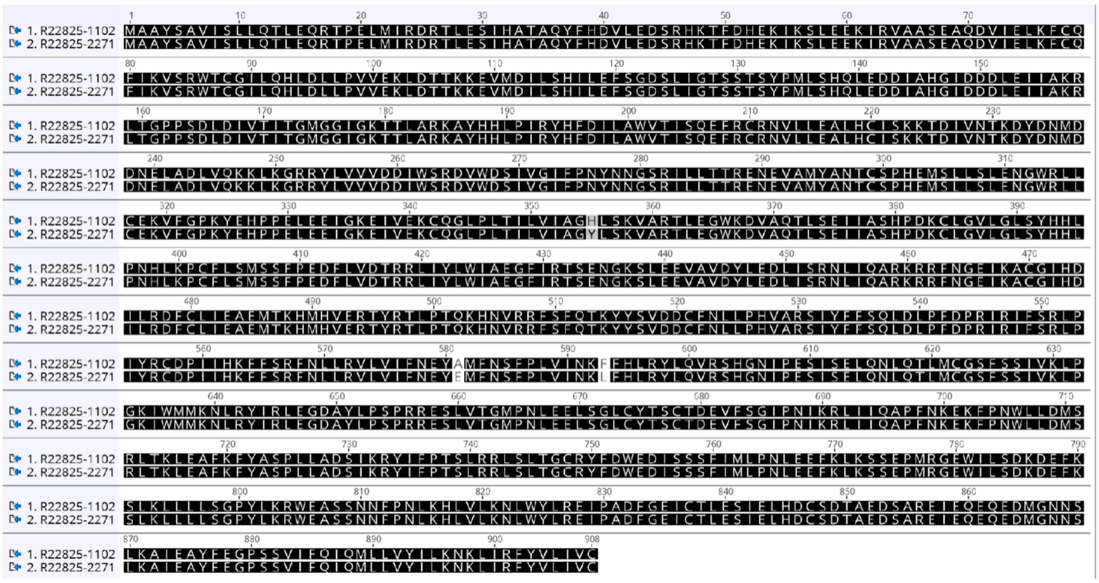


**Supplementary Figure 14**. Alignment of Rpi-amr4-1102 and Rpi-amr4-2271 proteins.

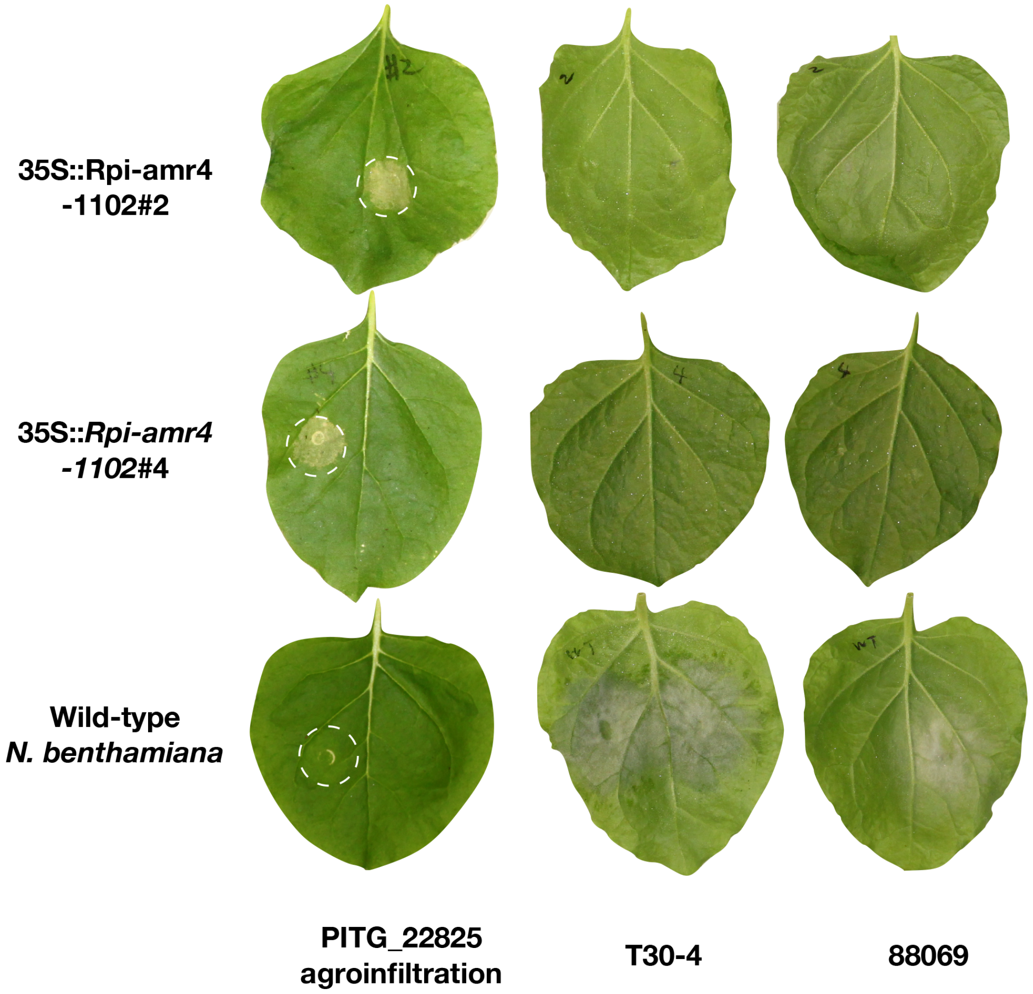


**Supplementary Figure 15**. PITG_22825 agroinfiltration and disease test on the *Rpi-amr4* transgenic *N. benthamiana*.

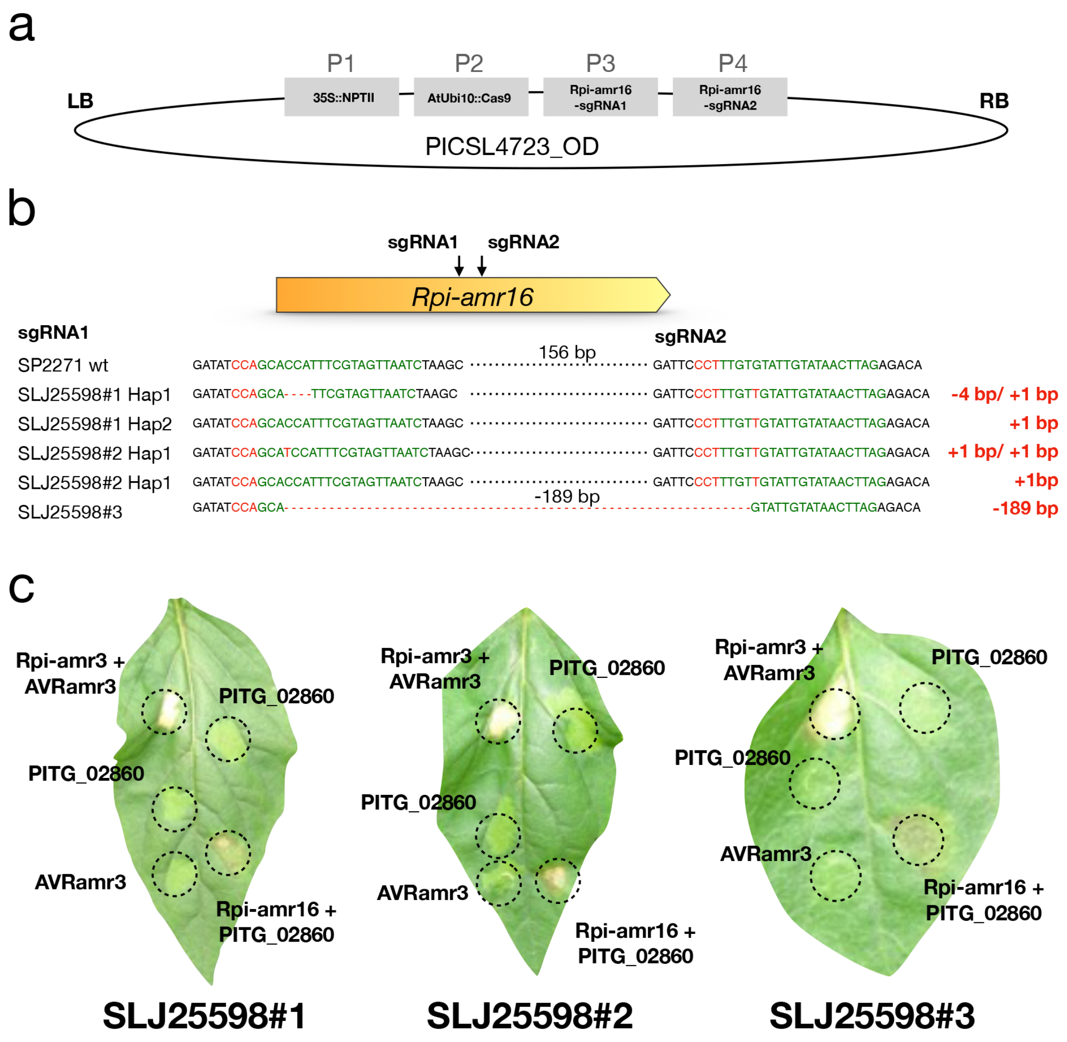


**Supplementary Figure 16**. *Rpi-amr16* knockout SP2271 lines. (a). The constructs used for the CRISPR-Cas9 knockout experiment; (b). Genotyping data of three *Rpi-amr16* lines SLJ25598#1, SLJ25598#2 and SLJ25598#3. (c). Phenotype of three selected knockout lines by agroinfiltration. Rpi-amr3 and AVRamr3 were used as controls.

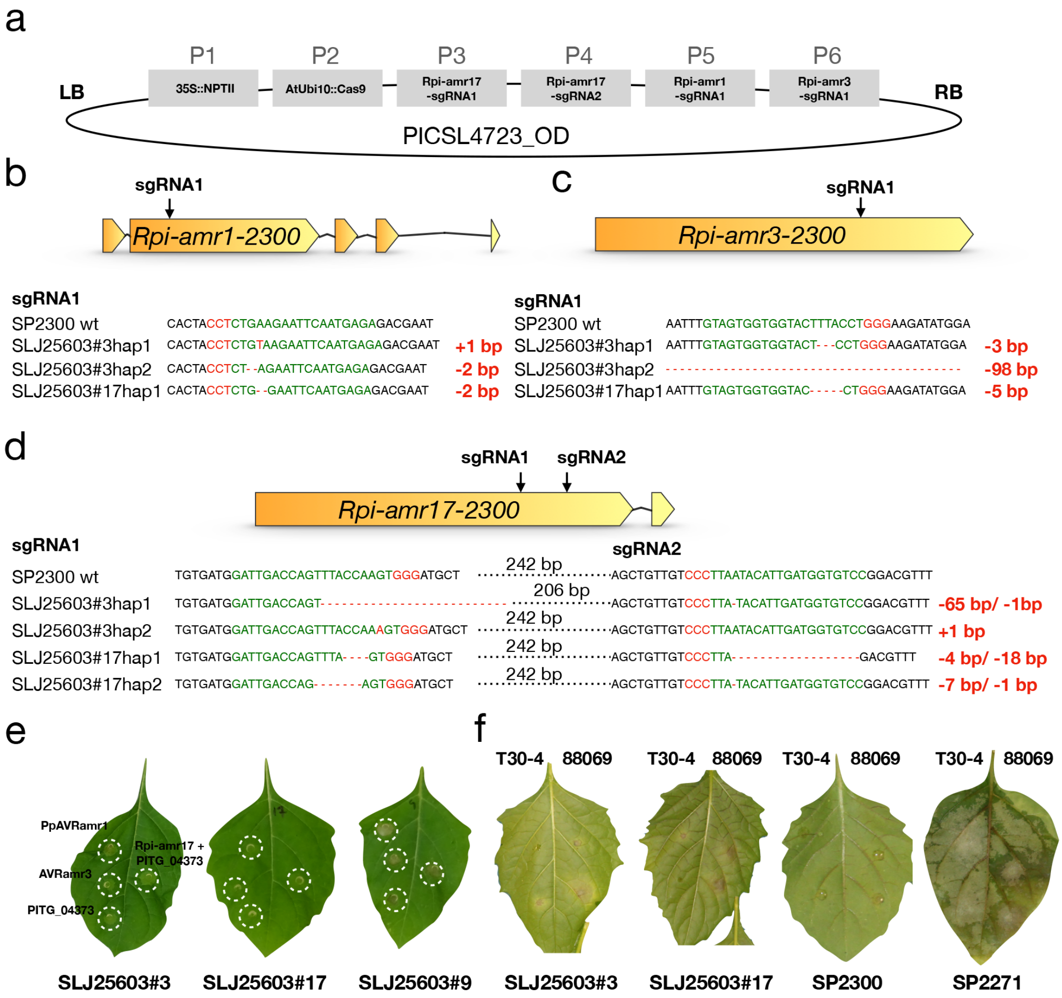


**Supplementary Figure 17**. *Rpi-amr1/Rpi-amr3/Rpi-amr17* triple knockout SP2300 lines. (a). The constructs used for the CRISPR-Cas9 knockout experiment; (b, c and d). Genotyping the *Rpi-amr1-2300* and *Rpi-amr3-2300* and *Rpi-amr17* in two knockout lines SLJ25603#3 and SLJ25603#17. (e). Phenotype of the triple knockout lines after expression of PpAVRamr1, AVRamr3 and PITG_04373. Co-expression of Rpi-amr17 and PITG_04373 was used as a control. (f). Disease test on the two triple knockout lines, *P. infestans* isolates T30-4 and 88069 was used in this assay. SP2300 and SP2271 were used as controls.
